# Circadian regulation of dentate gyrus excitability mediated by G-protein signaling

**DOI:** 10.1101/2022.06.28.497975

**Authors:** Jose Carlos Gonzalez, Haeun Lee, Angela M Vincent, Lacy K Goode, Gwendalyn D King, Karen L Gamble, Jacques I Wadiche, Linda Overstreet-Wadiche

## Abstract

The central circadian regulator within the suprachiasmatic nucleus transmits time of day information by a diurnal spiking rhythm that is driven by intrinsic activity of molecular clock genes controlling membrane excitability. Most brain regions, including the hippocampus, harbor similar intrinsic circadian transcriptional machinery but whether these molecular programs generate oscillations of membrane properties is unexplored. Here, we show that intrinsic excitability of mouse dentate granule neurons exhibits a 24-hour oscillation that controls spiking probability. Diurnal changes in excitability are mediated by antiphase G-protein regulation of potassium and sodium conductances to reduce excitability during the light phase. Disruption of the circadian transcriptional machinery by conditional deletion of *Bmal1* enhances excitability selectively during the light phase, increasing engram recruitment and spatial discrimination memory. These results reveal that circadian transcriptional machinery regulates intrinsic excitability, highlighting the role of cell-autonomous oscillations in hippocampal function and behavior.

**Highlights:** Dentate neurons exhibit a 24-hour oscillation of excitability with low excitability during the Light phase

Reduced excitability results from G-protein regulation of passive and active properties

Bmal1 deletion disrupts G-protein regulation to enhance excitability during the Light

Bmal1 deletion enhances the size of memory engrams and spatial discrimination

## Introduction

In mammals, the suprachiasmatic nucleus of the hypothalamus (SCN) is the primary pacemaker that governs daily rhythms by self-sustaining transcriptional-translational feedback loops that define a molecular clock (Hastings et al., 2018; Takahashi, 2017). These loops control the membrane excitability of SCN neurons, allowing it to transmit time of day information by an endogenous oscillation of characteristic spiking activity (Paul et al., 2020). Similar transcriptional loops drive local circadian programs of clock-controlled output genes in the hippocampus that interact with sleep-wake cycles to drive daily regulation of the synaptic proteome (Bruning et al., 2019; Noya et al., 2019; Zhang et al., 2014). While diurnal rhythms of synaptic function related to sleep-wake cycles are well-established (Bridi et al., 2020; Liu et al., 2010; Vyazovskiy et al., 2008), it is unclear whether local transcriptional clocks in the hippocampus also affect membrane excitability. Addressing this question is important for understanding how disruption of clock components contribute to abnormal excitability and epilepsy, as well as the daily rhythms of normal behavior (Debski et al., 2020; Snider et al., 2018; Wu et al., 2021; Zhang et al., 2021).

A 24-hr fluctuation of cortical-evoked population spikes and EPSPs in the dentate gyrus (DG) of awake rats and squirrel monkeys provided early evidence for circadian regulation of hippocampal input-output transformations (Barnes et al., 1977). The DG serves as a “gate” that filters incoming information from the entorhinal cortex to enable efficient information processing in CA3. The DG gate is maintained by the low propensity of the principal granule cells (GCs) to fire action potentials by a combination of synaptic and intrinsic mechanisms. GCs suppress propagation of neural activity through the cortico-hippocampal circuit by strong synaptic inhibition from local GABAergic interneurons (Coulter and Carlson, 2007; Dieni et al., 2013). Excitatory synaptic input related to dendritic complexity and differences in intrinsic excitability also contribute to the selection of GCs that comprise the active ensemble (Diamantaki et al., 2016; Erwin et al., 2020; Zhang et al., 2020). Thus, synaptic or intrinsic mechanisms could underlie time of day regulation of DG input-output transformations.

One intrinsic property that controls excitability is the steady-state membrane conductance that determines the resting membrane potential (RMP) and basal input resistance (IR). This resting conductance is primarily set by leak K^+^ and Na^+^ channels, with a ratio that varies in a cell-type specific manner. In GCs, G-protein coupled inwardly rectifying (GIRK) and other leak K^+^ conductances provide low intrinsic excitability by hyperpolarization and low membrane resistance (Gonzalez et al., 2018; Kim et al., 2020; Yarishkin et al., 2014). In central pacemaker neurons of flies and mice, molecular-clock controlled modulation of resting Na^+^ and K^+^ conductances are responsible for rhythms of membrane excitability that generate endogenous oscillation of spiking activity (Cao and Nitabach, 2008; Doi et al., 2011; Flourakis et al., 2015; Kuhlman and McMahon, 2004; Paul et al., 2016). Here we show that G-protein signaling, a pathway with robust circadian transcription independent of sleep-wake cycles (Noya *et al.,* 2019), mediates diurnal regulation of intrinsic excitability, synaptic recruitment and cognitive function in the dentate gyrus.

## Results

### Reduced recruitment of GCs during the Light phase compared to Dark phase

To compare excitability across the diurnal cycle, we prepared ventral hippocampal slices during the light phase (Zeitgeber Time, ZT 5.5 or 11.5 where ZT 12 refers to lights off) and performed recordings between projected ZT 8-11 (referred to as “Light phase”) or ZT 14-17 (“Dark phase”). This schedule avoids potential photic interference when comparing cellular properties in isolated slices during the Light and Dark phases. We recorded from visually identified GCs and stimulated the perforant path near the crest to minimize direct stimulation of local interneurons (**Figure 1A**). This paradigm reliably generated EPSPs at the lowest stimulus intensity but failed to recruit spikes during the Light phase, whereas up to ~40% of GCs were recruited to spike during the Dark phase (**Figure 1B**). We obtained similar results in GCs from different lines of transgenic mice (**Figure S1**). Thus, perforant path recruitment of GC spiking is robustly different across diurnal phase, with lower excitability during the Light phase.

**Figure 1.**
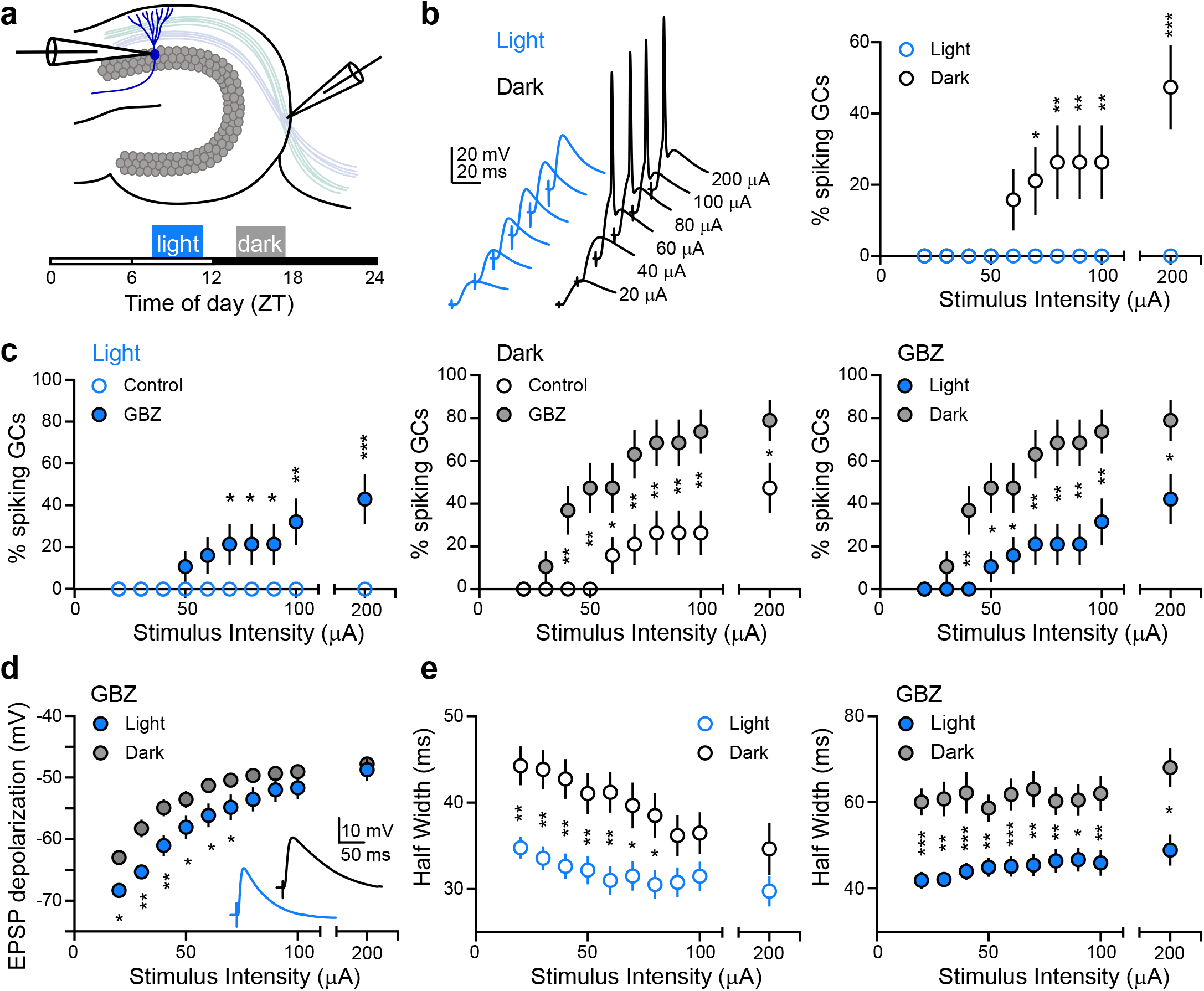
Reduced recruitment of GCs during the Light phase compared to Dark phase. (A) Current-clamp recordings (I=0) in response to stimulation of the perforant path were compared in two recording windows: ZT 8-11 (Light) and ZT 14-17 (Dark). (B) Left, example EPSPs at increasing stimulus intensities. Right, the percentage of GCs recruited to spike was reduced during the Light phase. (C) Gabazine (GBZ; 10 mM) increased spiking during both the Light (left) and Dark (middle) phases. In GBZ, a smaller percentage of GCs were recruited during the Light (right). (B-C) χ^2^ tests, *p<0.05; **p<0.01; ***p<0.001. (D) Depolarization by subthreshold EPSPs was reduced during the Light (in GBZ). Inset, EPSPs evoked at 100 mA. (E) EPSP half-widths were reduced during the Light in control (left) and GBZ (right). (D-E) Multiple comparison test *p<0.05; **p<0.01; ***p<0.001. (B-E) n = 19, 19. Symbols represent mean ± s.e.m.

GABA_A_ receptor-mediated inhibition strongly suppresses GC spiking (Dieni *et al.,* 2013; Yu et al., 2013). Accordingly, gabazine (GBZ; 10 μM) increased the percent of spiking GCs during both the Light and Dark phases (**Figure 1C**, left, middle). But in GBZ, the percentage of GCs that spiked remained lower during the Light phase compared with the Dark phase (**Figure 1C**, right). Thus, diurnal changes in GABAergic inhibition cannot account for differential spiking probabilities. To address other mechanisms, we compared subthreshold EPSPs in GBZ. At low stimulus intensities, the maximum depolarization was lower during the Light than Dark and the difference vanished at higher stimulus intensities as threshold was reached, suggesting no difference in spike threshold (**Figure 1D**). There was also no difference in the amplitude and rise time of subthreshold EPSPs when measured from respective resting potentials (**Figure S1**). However, there was a pronounced difference in the decay phase of subthreshold EPSPs. The half-width of EPSPs, both with inhibition intact and blocked, was shorter during the Light phase (**Figure 1E**). Since the membrane time constant (TC) dictates the decay phase of EPSPs, these results suggest that differences in GC spiking probability across Light and Dark phases could result from changes in intrinsic excitability.

### Reduced intrinsic excitability of GCs during the Light phase

To investigate whether the light period affects GC excitability through changes in intrinsic membrane properties, we bypassed synaptic stimulation and elicited action potential firing by current injections. During the Light phase, GCs exhibited less propensity for spiking, illustrated by a rightward shift in the spike-current relationship compared to the Dark phase (**Figure 2A**). There was also a significant reduction in passive excitability during the Light phase, with a more hyperpolarized RMP (−75.5 ± 0.3 vs −70.6 ± 0.5 mV), lower IR (281 ± 8 vs 345 ± 13 MΩ), and faster membrane time constant (27.8 ± 0.7 vs 34.2 ± 1.1 ms), culminating in higher rheobase, the current required to achieve threshold (66.1 ± 1.9 vs 50.7 ± 2.2 pA) (**Figure 2B**, violin plots). To assess whether these parameters varied systematically across time of day, we pooled recordings in 1-hour bins across 24 hours (**Figure 2B**, dot plots). Cosinor analysis reflected significant rhythmicity of RMP, IR, TC and rheobase (R^2^ = 0.37; 0.17; 0.28 and 0.17 respectively) and mesor, amplitude and acrophase values were evaluated for each parameter **(Table S1)**. The peak of GC excitability occurred at ZT 16 with the trough at ZT 4 (defined by acrophase values for rheobase) (**Figure 2C**).

**Figure 2.**
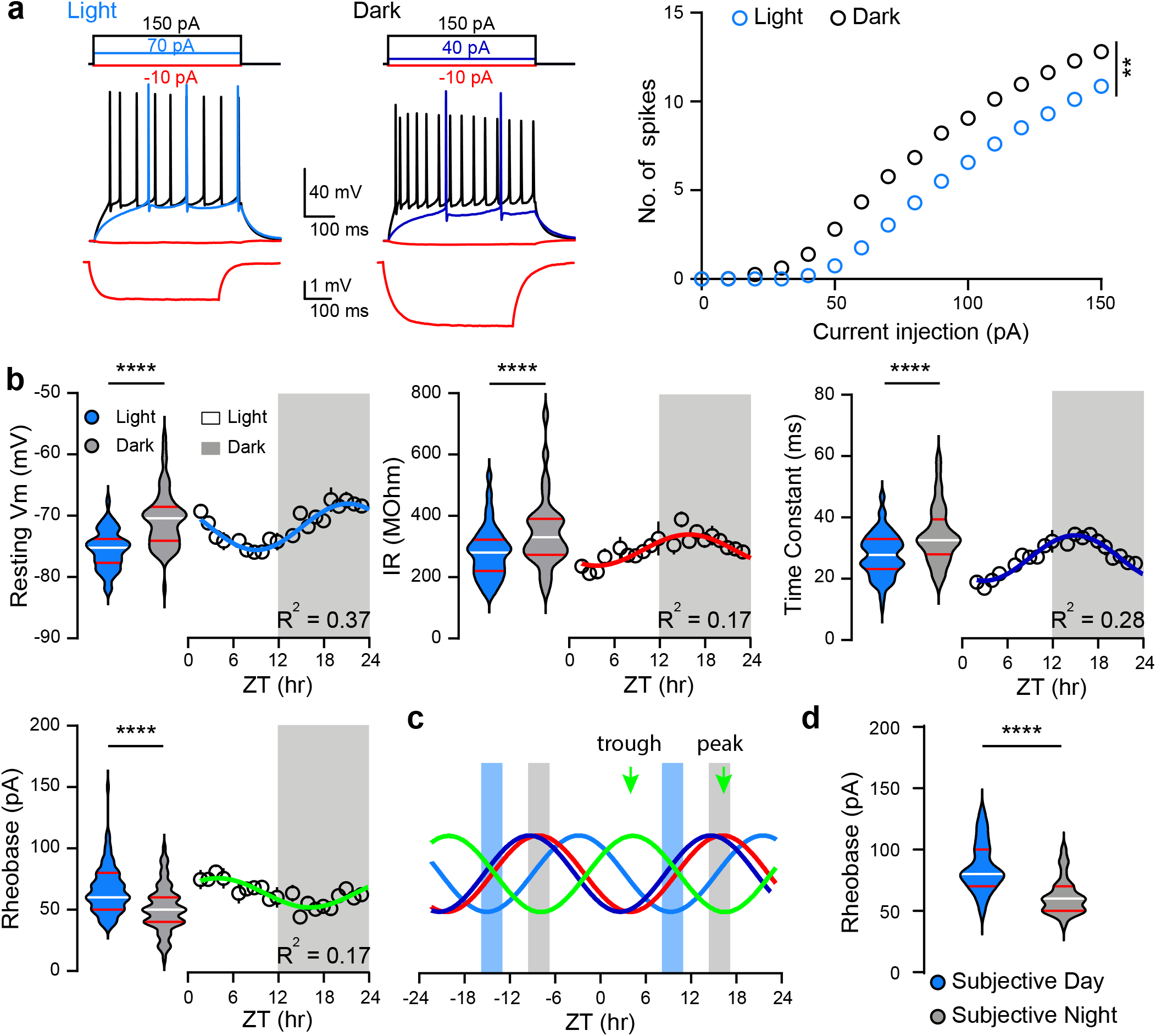
Reduced intrinsic excitability of GCs during the Light phase. (A) Left, examples of current injections (top) used to measure GC intrinsic properties (bottom). Right, number of spikes elicited by current steps revealed reduced spiking during the Light phase. n = 90 (Light); 69 (Dark). Unpaired t-test to compare area under the curve (AUC) t = 2.9 **p<0.01. (B) Violin plots show RMP, IR, TC and rheobase during the Light (blue) and Dark (gray) phase. Mann-Whitney U test, U = 1141 (RMP), U = 1997 (IR), U = 1845 (TC) and unpaired t test, t = 5.1 (rheobase). ****p<0,0001. Dot plots show intrinsic properties averaged in 1-hour bins fit with Cosinor function across 24 hours. Lights on at ZT0 and off at ZT12. n = 9-39. (C) Overlay of Cosinor fits for intrinsic properties, normalized in amplitude (same color code as in B). Blue and gray bars denote the Light and Dark recording windows, respectively. Arrows point to trough and peak of excitability based on acrophase for rheobase. (D) Violin plots show rheobase recorded at Subjective Day (Circadian Time or CT 8-11, where CT 12 refers to activity onset) and Subjective Night (CT 14-17) from mice housed in constant dark. Unpaired t-test, t = 5 ****p<0.0001. n = 27, 27. Bars on violin plots represent quartile and median values. Symbols represent mean ± s.e.m.

Since the light-dark cycle entrains many physiological and behavioral rhythms (LeGates et al., 2014), we also measured GC intrinsic excitability in mice kept in constant darkness (DD). After 30 days of DD, we made acute slices from mice during their Subjective Night (active) or Subjective Day (inactive) phases and found that GCs still exhibited differential intrinsic excitability similar to the 12:12 L-D values. This was exemplified by rheobase (85.5 ± 3.8 pA for Subjective Day vs 61.8 ± 2.7 pA for Subjective Night; **Figure 2D**), as well as other measures of intrinsic excitability, including the number of action potentials elicited by current injection (**Figure S2A**). Furthermore, GCs from both male and female mice showed reduced excitability during the Light and there was no interaction between sex and the recording window (**Figure S2B and Table S2**). These data show a daily rhythm of GC intrinsic excitability that likely contributes to diurnal regulation of DG excitability *in vivo* (Barnes *et al.,* 1977).

### Diurnal regulation of GC excitability requires GTP

Constitutive G-protein signaling contributes to low intrinsic excitability of mature dentate GCs (Gonzalez *et al.,* 2018), so we next tested whether diurnal differences in synaptic and intrinsic excitability relies on G-protein signaling by repeating recordings in the absence of Na-GTP (GTP^−^) in the intracellular solution. In this condition, there was no difference in spike recruitment by perforant path stimulation during the Light and Dark phases (**Figure 3A**), in contrast to what was shown in Figure 1C. Likewise, the number of APs generated by current injection was equalized during the Light and Dark phases (**Figure 3B**), and the intrinsic properties were also the same (RMP = −70.7 ± 0.2 vs −69.4 ± 0.7 mV; IR = 384 ± 18 vs 356 ± 21 MΩ; rheobase = 55.7 ± 3 vs 53.5 ± 3 pA). These results suggest that G-protein signaling is essential to generate diurnal rhythmicity and its absence during the Light phase phenocopies the higher excitability observed during the Dark. Indeed, the lack of GTP in the pipette increases excitability of GCs during the Light phase (see also (Gonzalez *et al.,* 2018)), whereas it had no effect on intrinsic excitability in GCs recorded during the Dark phase (**Figure S3**). Furthermore, diurnal regulation of excitability did not dependent of tonic GABA_A_ receptor activation, since GBZ had no effect on the intrinsic properties (**Figure S3**).

**Figure 3.**
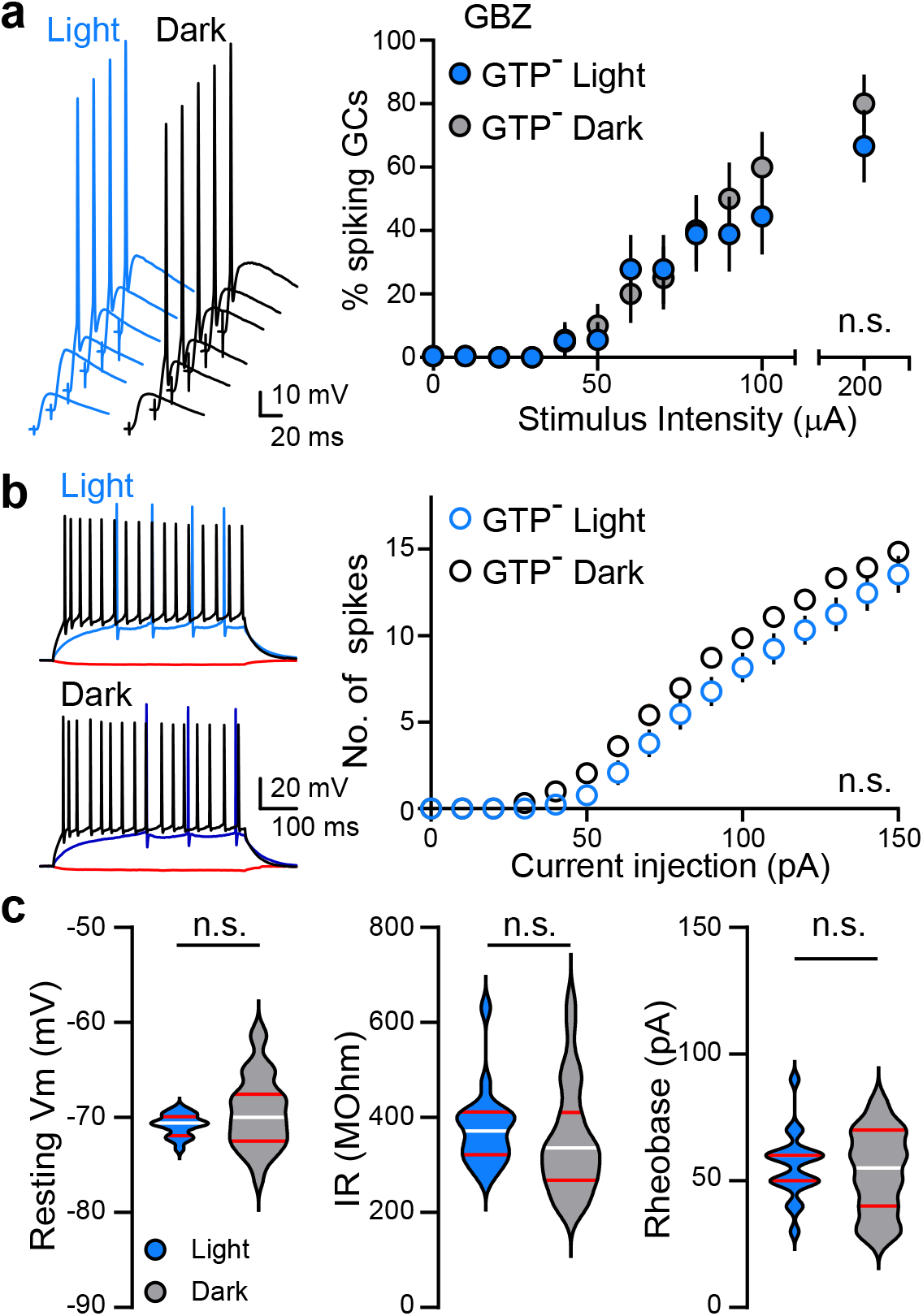
Diurnal regulation of GC excitability requires intracellular GTP. (A) Left, example EPSPs at increasing stimulus intensities in the absence of Na-GTP (GTP^−^) in the recording pipette. Right, the percentage of GCs recruited to spike by perforant path stimulation was similar during the Light and Dark phase; χ^2^ tests p > 0.05 at all stimulus intensities. (B) Left, example voltage traces in response to somatic current injections of −10 pA (red), rheobase (blue) and 150 pA (black). Right, the number of spikes elicited by current steps was similar during the Light and Dark phases. Unpaired t-test to compare area under the curve (AUC) t = 0.78 p=0.44. (C) Intrinsic properties of GCs recorded in the absence of GTP were similar between Light and Dark phases for RMP (left; Mann-Whitney U test, U = 212, p = 0.24), IR (middle; unpaired t test, t = 0.94, p = 0.35), and rheobase (right; unpaired t test, t = 0.5, p = 0.61). (A-C) n = 19 Light and 28 Dark. Bars on violin plots represent quartile and median values. Symbols represent mean ± s.e.m.

### Diurnal regulation of G-protein gated K^+^ and Na^+^ conductances

To address the role of G-protein regulated conductances to excitability, first, we tested the prediction that there is a larger GIRK-mediated conductance during the Light phase compared to the Dark phase, using a GABA_B_ inverse agonist (CGP55845, 10 μM; CGP) (Gonzalez *et al.,* 2018). As expected, CGP increased GC excitability, shifting the spike-current relationship to the left, depolarizing the RMP and increasing IR, thus reducing rheobase. However, CGP had no effect during the Dark phase (**Figure 4A,B**, red symbols). We also confirmed that differential effects of CGP were detected in Subjective Day and Subjective Night in DD mice (**Figure S4A,C**). To determine whether the differential effect of CGP is mediated via regulation of constitutive activity or down-regulation of GIRK channels expressed on the membrane, we compared the actions of the GIRK-selective activator ML297 (10 μM) in the Light and Dark phase. ML297 reduced the number of action potentials elicited by current injection, hyperpolarized the RMP and decreased IR and excitability in both Light and Dark phases (**Figure 4A,B**, green symbols). In the presence of ML297, there were no differences in excitability between Light and Dark phases (**Figure S4B,D**). Together, these results suggest that regulation of GIRK channel activity, rather than changes in the number of GIRK channels, contributes to the GTP-dependent diurnal rhythms in GC excitability.

**Figure 4.**
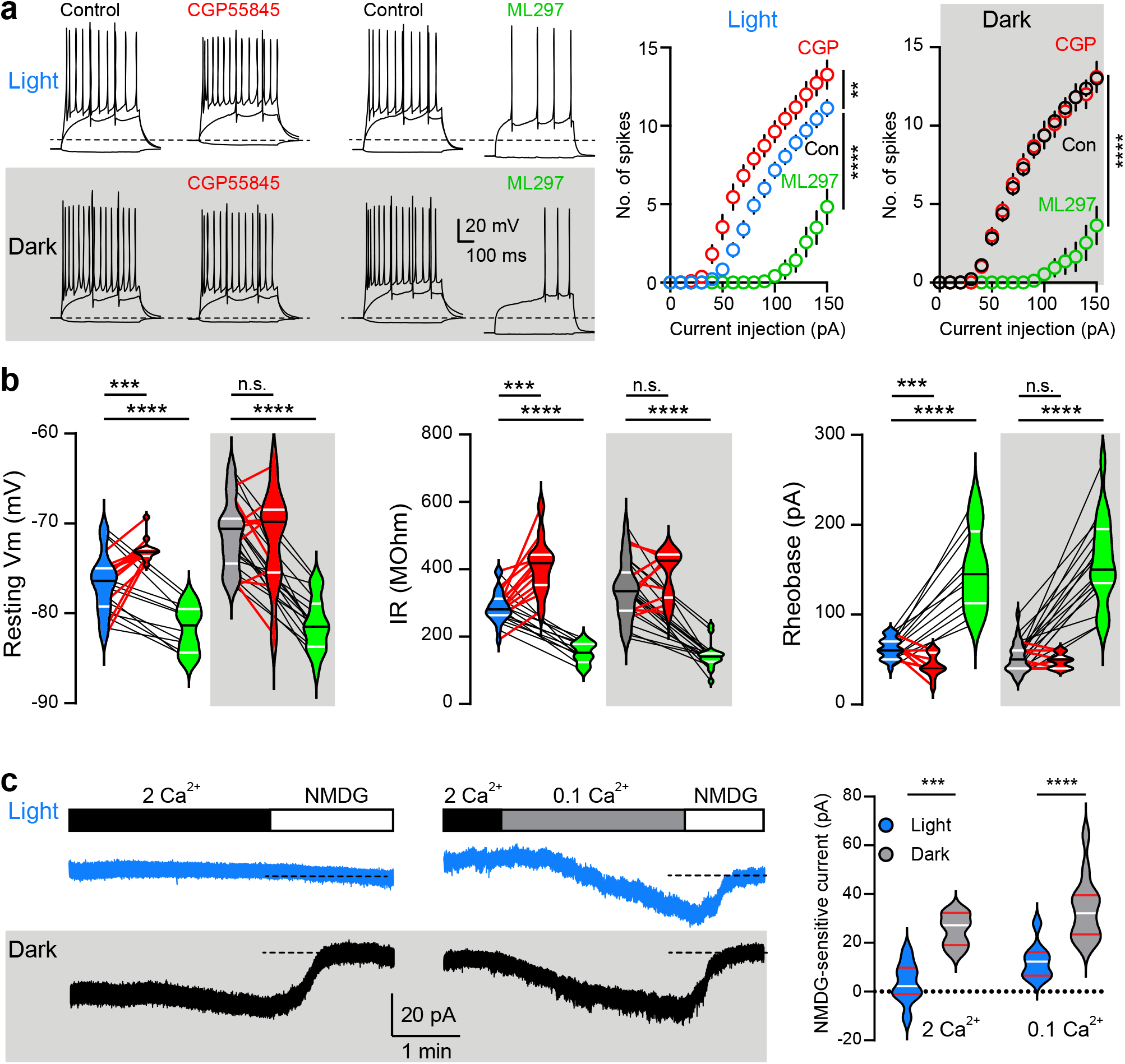
Diurnal regulation of G-protein gated K^+^ and Na^+^ conductances. (A) Left, example voltage traces in response to current injections (−10 pA, rheobase and +150 pA) in CGP55845 (10 mM; red) or ML297 (10 mM; green) during Light (top) or Dark phases (bottom). Dotted line indicates −70 mV. Right, the number of action potentials elicited by current injections in the same conditions. Note that controls for application of CGP and ML297 (a paired analysis) were pooled. In Light, n = 11 for CGP, n = 12 for ML297. In Dark, n = 10 for CGP, n = 17 for ML297. One-way ANOVA to compare area under the curve (AUC) F_(2,43)_ = 46.9 and F_(2,51)_ = 65.2 for Light and Dark respectively followed by Tukey’s multiple comparisons test. **p<0.01; ****p<0.0001. (B) Violin plots showing effects of CGP55845 (red) or ML297 (green) during the Light and Dark phases. For RMP in Light, CGP paired t-test t = 5.6; ML297 t = 7.1. In Dark, CGP t = 0.2; ML297 t = 10.3. For IR in Light, CGP paired t-test t = 6.1; ML297 t = 8.9. In Dark, CGP t = 1.7; ML297 t = 8.7. For Rheobase in Light, CGP paired t-test t = 5.5; ML297 t = 8.1. In Dark, CGP t = 0.0; ML297 t = 9.7. n.s. p > 0.05; *** p < 0.001; **** p < 0.0001. (C) Left, examples of pharmacologically-isolated Na^+^ currents voltage-clamped at −70 mV during the Light (blue) or Dark (gray shadow) phases. Dashed line indicates the current blocked by NMDG in 2 mM extracellular Ca^2+^ (left) or 0.1 mM Ca^2+^ (right). Right, violin plots show larger NMDG-sensitive Na+ currents during the Dark phase at both [Ca^2+^]s. Dashed line at 0. Unpaired t-test in 2 mM Ca^2+^, t = 5.6; in 0.1 mM Ca^2+^, t = 4.5. *** p < 0.001; **** p < 0.0001. n = 9, 7, 9 and 13 for Light 2 mM Ca^2+^, Dark 2 mM Ca^2+^, Light 0.1 mM Ca^2+^ and Dark 0.1 mM Ca^2+^ respectively. Bars on violin plots represent quartile and median values. Symbols represent mean ± s.e.m.

Second, we evaluated the role of non-selective Na^+^ leak channel (NALCN) activity that is inhibited by G-protein signaling (Lu et al., 2007; Philippart and Khaliq, 2018). We isolated TTX-insensitive Na^+^ conductance in the presence of synaptic and K^+^ channel blockers and measured the current sensitive to substitution of Na^+^ ions by N-methyl-D-glutamine (NMDG). During the Light phase, there was no significant shift in membrane current by Na^+^ replacement (3.5 ± 2.7 pA), suggesting that Na^+^ channels may be blocked by constitutive G-protein signaling. In contrast, during the Dark phase, NMDG caused an outward current (25.3 ± 2.5 pA) indicating the removal of an inward tonic Na^+^ current (**Figure 4C**). Repeating the experiment in 0.1 mM Ca^2+^, which potentiates NALCN channel activity(Lu et al., 2010), revealed a small persistent Na^+^ current during the Light phase (12.4 ± 2.4 pA) that was larger during the Dark phase (33.9 ± 3.5 pA). These results suggest that constitutive activity of βγ subunits during the day reduces GC intrinsic excitability by blocking NALCN channels that are active during the Dark phase. In other words, diurnal rhythms of GC excitability are mediated by antiphase regulation of G-protein dependent K^+^ and Na^+^ conductances, reminiscent of the antiphase cycles that drives circadian control of membrane potential in master clock neurons (Doi *et al.,* 2011; Flourakis *et al.,* 2015; Paul *et al.,* 2016).

### GTP-dependent diurnal regulation of spike bursts

GCs *in vivo* can fire brief high-frequency bursts of 2-4 action potentials important for recruitment of CA3 neurons via strongly facilitating mossy fiber output synapses (Pernia-Andrade and Jonas, 2014). Although there were no significant differences in the properties of individual action potentials, including threshold, amplitude and width, across the light cycle (not shown), we observed a striking diurnal difference in the propensity for burst firing that was dependent upon intact G-protein signaling.

During the Light phase, GCs exclusively exhibited a single action potential in response to rheobase current injection (**Figure 5A**) and perforant path stimulation (**Figure 5B**). However, GCs recorded in the Dark phase or in absence of intracellular GTP during the Light phase often exhibited a burst of action potentials, defined by 2 or 3 action potentials at frequencies over 100 Hz. At rheobase, 31% and 36% of GCs showed bursting during the Dark phase or in the absence of GTP during the Light phase, respectively (**Figure 5A**). Likewise, no GCs displayed action potential bursts in response to perforant path stimulation during the Light phase, yet 42% of GCs in the Dark phase exhibted bursting, and when GTP was excluded from the recording pipette, 18% of GCs in the Light phase exhibited bursting (**Figure 5B**). These results suggest that additional mechanisms beyond passive GC properties by constitutive K^+^ and Na^+^ conductances likely contribute to reduced excitability during the Light phase. One likely candidate is T-type Ca^2+^ channels that are implicated in GC bursting and a well-known target for suppression by Gi/o signaling (Dumenieu et al., 2018).

**Figure 5.**
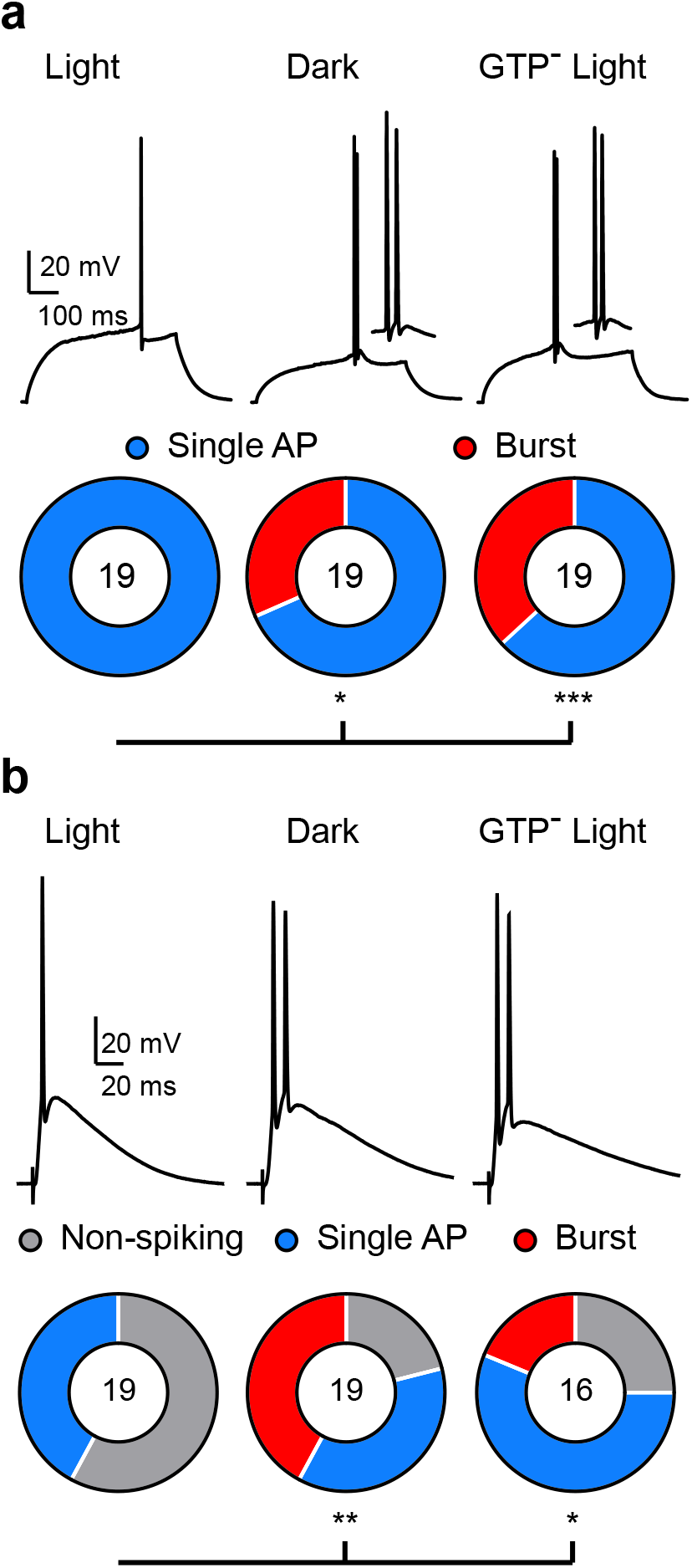
GTP-dependent diurnal regulation of spike burst generation. (A) Example voltage traces in response to rheobase current injection during the Light phase (left), Dark phase (middle) and Light phase using a GTP^−^ internal (right). Insets show enlargment of the action potentials (APs). Sectors represent the fraction of GCs with single or multiple APs in each condition (χ^2^ tests, *p<0.05; ***p<0.001). (B) Examples of APs evoked in response to perforant path stimulation at 200 mA. Sectors represent the percentage of GCs with each spike pattern, as well as non-spiking cells. χ^2^ tests, *p<0.05; **p<0.01. No. of cells is indicated.

### Diurnal regulation of synaptic recruitment is disrupted by *Bmal1* cKO

Based on the daily cycle of core clock gene expression in hippocampus (Jilg et al., 2010), we wondered whether diurnal regulation of GC excitability relies on a cell-autonomous transcriptional oscillation. Thus, we generated SCN-independent disruption of the molecular clock by conditional *Bmal1* ablation (Bunger et al., 2000), targeting GCs using *Pomc-Cre* with visualization by the Ai14 reporter (tdTomato; tdT). *Pomc-Cre* targets a majority of postmitotic dentate GCs with sparse expression in other brain regions (McHugh et al., 2007; Adlaf et al., 2017). Consistent with a restricted expression pattern, *Pomc-Cre:tdT:Bmal1^lox/lox^* mice (Bmal1 conditional KO; cKO) exhibited normal circadian locomotor rhythms in both LD and DD conditions, indicating intact SCN function (**Figure S6**).

In cKO mice, BMAL1 immunolabeling revealed that most tdT^+^ GCs have low BMAL1 immunoreactivity whereas tdT^−^ GCs located in the outer region of the granule cell layer express stronger BMAL1 immunoreactivity (**Figure 6A**). To verify conditional deletion, we measured the intensity of BMAL1 immunoreactivity in tdT^+^ GCs from *Pomc-Cre:tdT:Bmal1^lox/lox^* (cKO) and *Pomc-Cre:tdT:Bmal1^wt/wt^* (control) mice and found a negative correlation only in cKO mice (**Figure 6B**). The presence of Cre positive and negative GCs allowed us to determine the cell-autonomous effect of *Bmal1* cKO by recording from neighboring tdT^+^ GCs (*Bmal1* cKO) and tdT^−^ (*Bmal1* WT) within the same slice, and to directly compare synaptic recruitment by the same afferent fiber stimulation (**Figure 6C**) (Adlaf *et al.,* 2017). During the Light phase, *Bmal1* cKO GCs were more likely to spike in response to perforant path stimulation compared to neighboring WT GCs, revealing a cell-autonomous increase in afferent recruitment at most stimulus intensities (**Figure 6D**, left). However, there was no difference in perforant path-evoked spiking during the Dark phase (**Figure 6D**, right). Interestingly, we also found a burst-like pattern on the 28% of *Bmal1* cKO GCs during the Light phase (**Figure 6E**). These results reveal a cell-autonomous role of the molecular clock in reducing excitability of GCs specifically during the Light phase. The similar (high) excitability of tdT^+^ and tdT^−^ GCs during the Dark reinforces the hypothesis that diurnal changes in GC excitability result from a molecular clock-dependent reduced excitability during the Light phase, rather than increased excitability during the Dark phase.

**Figure 6.**
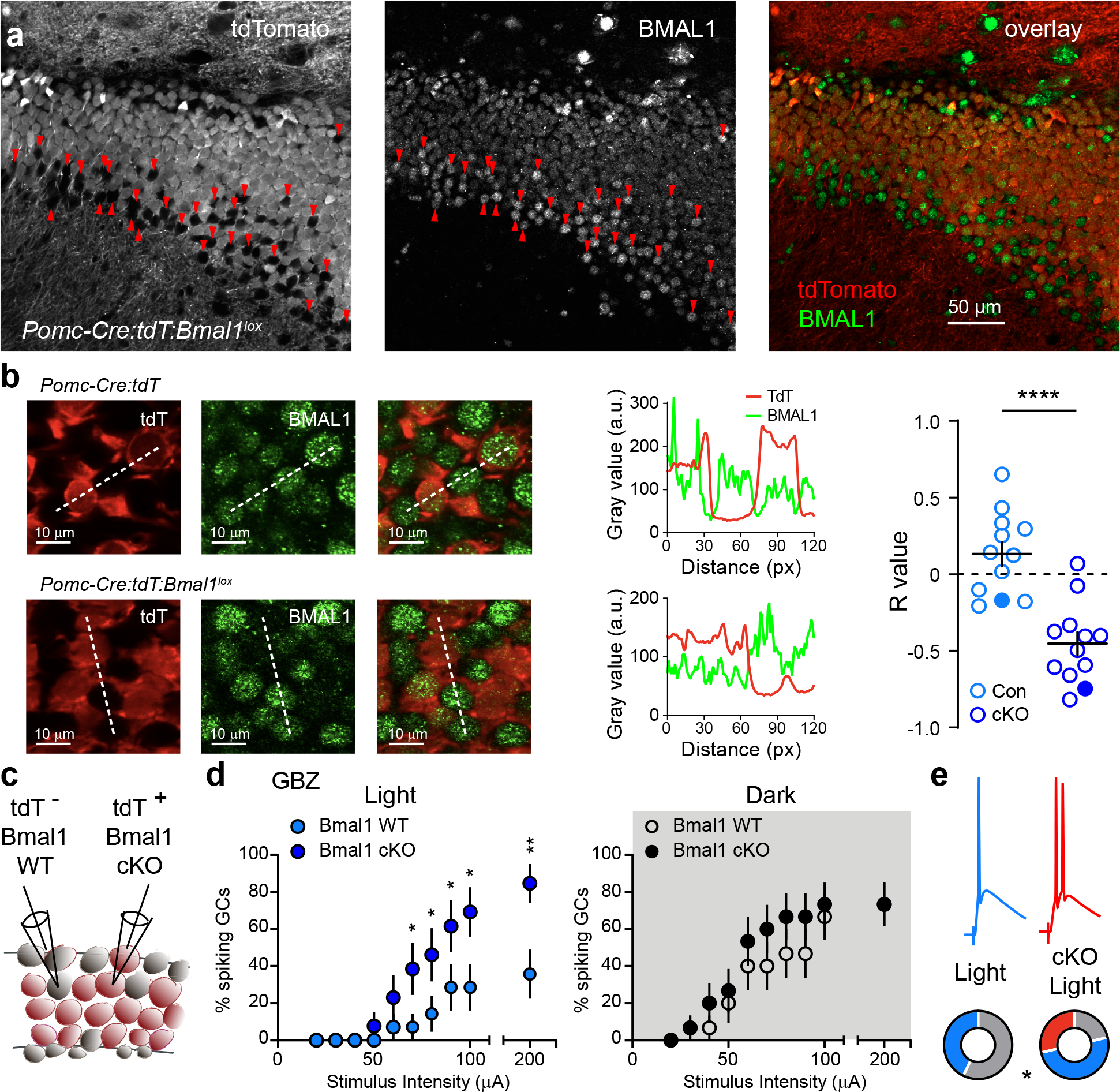
Diurnal regulation of synaptic recruitment is disrupted by *Bmal1* cKO. (A) Confocal images from a *Pomc-Cre:tdT:Bmal1^lox/lox^* (cKO) mouse showing tdTomato (left), BMAL1 (middle) and overlay (right). Arrows point to tdT^−^/BMAL1^+^ GCs. (B) Confocal images of tdT (left), BMAL1 (middle) and overlay (right) from *Pomc-Cre:tdT* (con; top) or cKO (bottom). Dashed line shows where fluorescence was registered in arbitrary units (a.u.). Examples of tdT (red) and BMAL1 (green) fluorescence from dotted lines (middle) with summary of R values (right). Filled symbols represent examples. Unpaired t-test t = 5.4. **** p < 0.0001 from 2 con and 6 cKo mice. (C) Neighboring tdT^−^/*Bmal1* WT and tdT^+^/*Bmal1* cKO GCs were recorded in the same slices. (D) The % of GCs that spike in response to perforant path stimulation was increased in *Bmal1* cKO GCs in the Light phase (left, n=14 pairs of GCs) but not in the Dark phase (right, n=15 pairs of GCs). (E) *Bmal1* cKO GCs exhibited burst firing in response to perforant path stimulation during Light phase. Sectors represent non-spiking (grey), single spikes (blue) and burst firing patterns (red). n = 14 pairs of GCs. (D-E) χ^2^ tests, *p<0.05; **p<0.01. Symbols represent mean ± s.e.m.

To validate regulation of GC excitability by the cell autonomous circadian clock, we directly compared GC intrinsic excitability and its regulation by G-protein signaling in *Bmal1* cKO GCs. *Bmal1* WT GCs (tdT^−^) exhibited the expected diurnal difference in intrinsic excitability shown in Figure 2, with a rightward shift in the spike-current relationship during the Light phase (**Figure 7A** open symbols) and single action potentials (**Figure 7B**), along with a hyperpolarized RMP (−75.7 ± 0.4 vs −69.1 ± 1 mV) and increased rheobase (80 ± 3.3 vs 60.6 ± 4.2 pA) compared to the Dark phase (**Figure 7C**, light blue versus gray violin plots). However, neighboring *Bmal1* cKO GCs (tdT+) failed to exhibit a diurnal difference in intrinsic excitability. Instead, during the Light phase *Bmal1* cKO GCs phenocopied the Dark phase excitability (**Figure 7A**, solid symbols) and action potential burst pattern (**Figure 7B**), with the same depolarized RMP (−68.3 ± 0.8 vs −70 ± 0.8 mV) and rheobase (63.6 ± 4 vs 64.6 ± 4.4 pA) as during the Dark phase (**Fig. 7c**, dark blue and black violin plots). Thus, *Bmal1* cKO GCs were more excitable than neighboring WT GCs only during the Light phase (**Figure 7A-C**, light blue open symbols vs dark blue solid symbols and violin plots). Importantly, the lack of diurnal regulation of synaptic recruitment and intrinsic excitability were dependent on the *Bmal1* floxed gene rather than Cre expression, as comparing intrinsic excitability and synaptic recruitment in tdT^+^ and tdT^−^ GCs from *Pomc-Cre:tdT:Bmal1^wt/wt^* mice revealed no differences (**Figure S7**). Thus, deletion of *Bmal1* leads to a cell-autonomous deficit in diurnal regulation of GC excitability by preventing reduced excitability during the Light phase.

**Figure 7.**
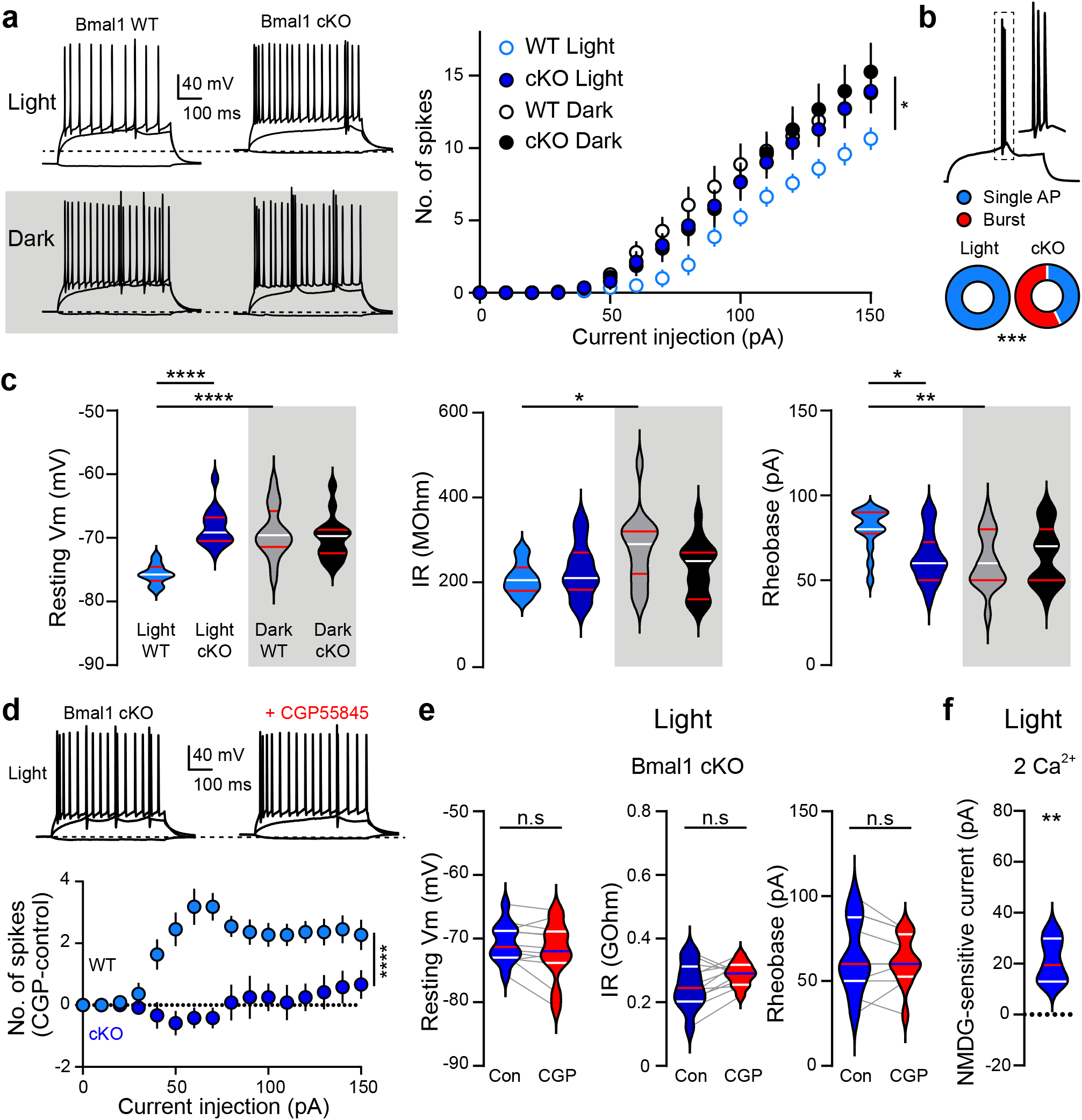
Diurnal regulation of synaptic recruitment is disrupted by *Bmal1* cKO. (A) Left, examples of voltage traces in response to current injections (of −10 pA, rheobase and 150 pA) for tdT^−^ (Bmal1 WT) and tdT^+^ (Bmal1 cKO) GCs from cKO mice. Dashed line marks avg RMP during the Dark phase. Right, number of APs elicited by current injections in the same conditions, blue symbols denote Light phase and black denotes Dark phase. Welch’s ANOVA to compare area under the curve (AUC) F_(3,28)_ = 3.44 followed by Dunnett’s multiple comparisons test. *p<0.05. (B) Representative bursts recorded at rheobase from *Bmal1* cKO GCs during the Light phase. Sectors represent differences in discharge profile. χ^2^ tests, ***p<0.001. n = 14 pairs of GCs. (C) Intrinsic properties of WT and cKO GCs during the Light (blue) and Dark (black) phases. One-way ANOVA for RMP (F_(3,54)_ = 16.24), IR (F_(3,54)_ = 3.06) and rheobase (F_(3,54)_ = 4.52) followed by Tukey’s multiple comparison test. * p < 0.05; ** p < 0.01; **** p < 0.0001. (A-C) Sample size n = 14 *Bmal1* WT and cKO GCs in the Light phase, n = 15 *Bmal1* WT and cKO GCs in the Dark phase. (D) Top, example voltage traces before and after application of GCP55845 (10 mM; CGP) in a cKO GC in the Light phase. Dashed line marks avg RMP during the Dark phase. Bottom, comparison of the number of action potentials elicited by CGP perfusion (CGP – control) in Bmal1 WT and *Bmal1* cKO GCs in the Light phase. Unpaired t-test to compare area under the curve (AUC) t = 5.1; **** p<0.0001. (E) Violin plots of intrinsic properties from *Bmal1* cKO GCs show that CGP had no effect during the Light phase. Paired t test for RMP (t = 1.9), IR (t = 2) and rheobase (t = 0.8); p > 0.05. n = 12. (F) Violin plot showing a robust NMDG-sensitive current in in *Bmal1* cKO GCs in the Light phase (contrast with Figure 4C). Unpaired t test t = 3.7. ** p < 0.01.n = 5. Bars on violin plots represent quartile and median values. Symbols represent mean ± s.e.m.

To confirm that the loss of diurnal regulation of excitability results from impairment of G-protein regulated K^+^ and Na^+^ conductances, we assessed these conductances in *Bmal1* cKO GCs during the Light phase. In contrast to WT GCs that exhibit enhanced excitability in response to CGP, *Bmal1* cKO GCs showed no changes in the spike-current relationship (**Figure 7D**) and intrinsic properties (−70.6 ± 0.9 vs - 71.9 ± 1.3 mV; 251 ± 20 vs 288 ± 10 MΩ; 64.1 ± 6.2 vs 61.5 ± 4.2 pA for RMP, IR and rheobase, respectively) (**Figure 7E**). Furthermore, *Bmal1* cKO GCs exhibited a pronounced constitutive Na^+^ conductance (21 ± 3.9 pA) that was absent in WT GCs (3.5 ± 2.7 pA, from Figure 4) (**Figure 7F**). Thus, *Bmal1* deletion results in a cell-autonomous loss of G-protein regulated K^+^ and Na^+^ conductances during the Light phase. Interestingly, while the spike-current relationship, RMP, rheobase and constitutive K^+^ and Na^+^ conductances in Bmal1 cKO GCs phenotypes the Dark phase, the IR did not (**Figure 7C**, middle). This suggests that the loss of the constitutive K^+^ conductance (expected to increase IR) is countered by an enhanced leak Na^+^ conductance (expected to decrease IR), that together mask changes in IR despite the robust consequences on excitability.

### Enhanced GC excitability in *Bmal1* cKO mice promotes engram formation

Intrinsic excitability of GCs controls the flux of spatial and sensory information through the DG and is thus poised to control the size of engram ensembles underlying contextual memories (Tonegawa et al., 2015; Zhang *et al.,* 2020). Hence, we tested whether the enhanced excitability of *Bmal1* cKO GCs during the Light phase translates to an increase in the size of a memory engram *in vivo*. Initial comparison of *Bmal1* cKO mice and littermate controls (in the Light phase) revealed no obvious differences in behavior with similar velocity and distance traveled in an open field (**Figure 8A**). To establish a contextual memory, we subjected mice to a standard fear conditioning protocol that generated a high level of freezing immediately after the shock, as well as when mice were reintroduced to the shock chamber 24 hours later (**Figure 8B**). One hour after re-instating the fear memory in the shock chamber, we detected an increase in the total number of c-Fos^+^ cells in *Bmal1* cKO (**Figure 8C**). As previously described, most c-Fos^+^ cells were in the outer layer of the suprapyramidal blade of the GCL, potentially representing semi-lunar GCs (Erwin *et al.,* 2020). In *Bmal1* cKO mice, the increase in c-Fos^+^ cells likewise occurred in the suprapyramidal blade with no detectable difference in the infrapyramidal blade. Furthermore, the increase in c-Fos^+^ cells was accounted for by an increase in the inner half of the GCL, consistent with the predominate location of *Pomc-Cre* targeted GCs (**Figure 8D**; see Figure 6). Thus, enhancing excitability of GCs by conditional deletion of *Bmal1* resulted in an increase in c-Fos expression *in vivo*, suggesting that diurnal variation of GC excitability under control of *Bmal1* contributes to the size of DG engrams.

**Figure 8.**
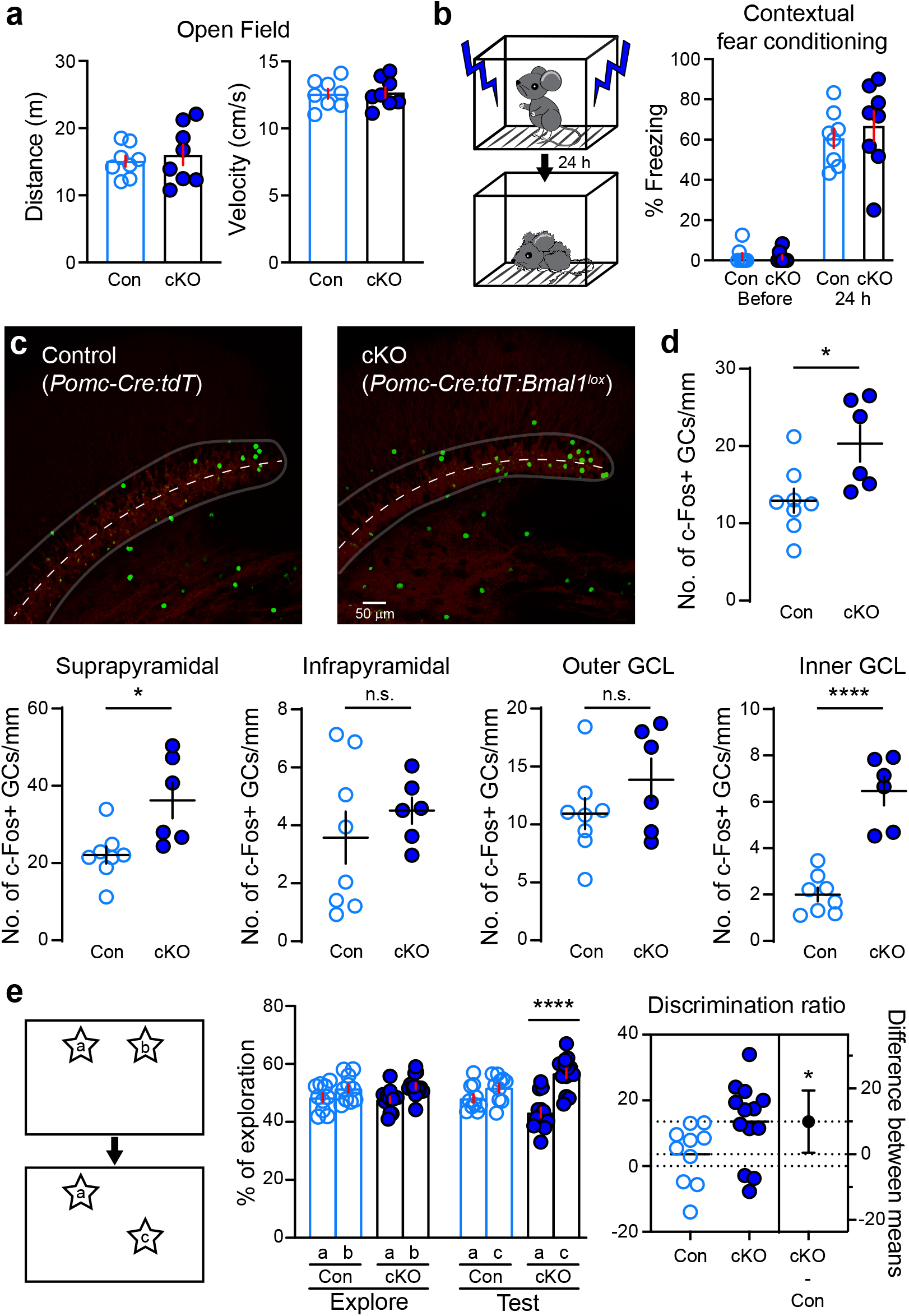
Enhanced GC excitability in *Bmal1* cKO mice promotes engram formation. (A) Open field total distance traveled and velocity for *Pomc-Cre:tdT* (Control; Con) and *Pomc-Cre:tdT:Bmal1^lox/lox^* (cKO) mice. (B) Left, depiction of fear conditioning protocol that was quantified (right) as the % of time spent freezing on training day prior to foot shock and 24 hours later when mice are returned to the same context. (C) Confocal images of c-Fos immunohistochemistry (green) and tdT expression (red) from Con and cKO mice. The granule cell layer (GCL) is outlined and the dashed line indicates the division used use quantify cells in the outer and inner GCL. (D) Quantification of c-Fos labeled GCs in the whole GCL (Unpaired t test, t = 2.7 p < 0.05), suprapyramidal blade (Unpaired t test, t = 2.9, * p < 0.05), infrapyramidal blade (Unpaired t test, t = 0.8, p > 0.05), outer GCL (Unpaired t test, t = 1.3, p > 0.05) and inner GCL (Unpaired t test, t = 7, *** p < 0.0001). n = 8, 6 for Con and cKO mice, respectively. (E) Left, depiction of object location memory (OLM) task. Middle, histograms represent % of time exploring the indicated object on day 1 (explore) and 24 hours later when one object was displaced (Test). One-way ANOVA followed by Sidak multiple comparison post hoc test (F_(7,84)_ = 8.9, p < 0.001). Right, estimation plot of the discrimination ratio. Unpaired t test, t = 2.1, * p < 0.05. n = 10 and n = 13 for Con and cKO respectively. Symbols represent mean ± s.e.m.

Although *Bmal1* cKO mice exhibited normal behavior in the fear conditioning protocol, we wondered whether changes in GC intrinsic properties could alter a behavioral task that is specifically dependent upon GC excitability (Pofahl et al., 2021). Object location memory (OLM) is a simple test of spatial memory that evaluates the ability to discriminate and recollect when an object is moved from its original location. It relies on the DG-CA3 circuit and exhibits robust diurnal variation with poor performance during the Light phase (Snider et al., 2016). During the training day of this task, *Bmal1* cKO and control mice spent a similar amount of time exploring two objects in an open field. However, when reintroduced to the field 24 hours later, only *Bmal1* cKO mice significantly increased the time spent exploring the object placed in the new location, resulting in an increase in the discrimination ratio for the moved object (**Figure 8E**). These results suggest that circadian regulation of GC excitability via G-protein signaling contributes to diurnal regulation of spatial discrimination memory, with reduced excitability during the Light phase contributing to impaired discrimination.

## Discussion

Circadian regulation of membrane physiology beyond the SCN is largely unexplored (reviewed in (Paul *et al.,* 2020)). Early evidence for diurnal modulation of excitability showed that DG population spikes in rats and squirrel monkeys are enhanced during active periods (Barnes *et al.,* 1977). However, the cellular mechanisms contributing to differential excitability and whether these oscillations are controlled by the central circadian clock or under cell-intrinsic control are not clear. In contrast, it is well known that SCN neurons, under the supervision of clock transcriptional components, exhibit circadian oscillations in membrane intrinsic properties that are regulated by various mechanisms including channel trafficking, kinase activity and auxiliary subunit expression (Farajnia et al., 2015; Flourakis *et al.,* 2015; Kuhlman and McMahon, 2004; Paul *et al.,* 2016; Whitt et al., 2016).

Here we demonstrate that GCs exhibit reduced excitability during the Light phase in a manner independent of GABA_A_ receptor mediated synaptic inhibition. Measures of intrinsic excitability including resting membrane potential, input resistance and rheobase oscillate with a period of approximately 24 hours controlled by a GTP-dependent mechanism. Constitutive antiphase K^+^ and Na^+^ leak conductances, both regulated by G_i/o_ signaling, drive rhythmic depolarization and hyperpolarization when *Bmal1* expression is intact. Hence, chronodisruption by conditional deletion of *Bmal1* enhanced GC excitability only during the Light phase. These results show that cell-autonomous transcriptional regulation requiring *Bmal1* reduces GC excitability during the Light phase via disruption of G protein signaling, illustrating a novel mechanism coupling membrane excitability by antiphase K^+^ and Na^+^ conductances to the peripheral molecular clock.

The efficiency of synaptic recruitment depends on characteristics of the underlying synaptic conductances and the passive and active properties of the neuronal membrane. Here we found reduced cortical recruitment during the light phase. Low intrinsic excitability contributes to differential recruitment by synaptic stimulation (Figure 1) because 1) the observed differences were readily apparent using current injections to isolate membrane properties; 2) differences remained in the presence of GBZ but were 3) absent when we excluded GTP from the pipette recording solution; and 4) our conditional deletion model allows us to compare synaptic recruitment of neighboring GCs with intrinsic excitability suppressed or intact by *Bmal1* deletion. In addition to our focus on GC intrinsic excitability, it is likely that multiple EC-hippocampal network components exhibit diurnal regulation excitability including the network of GABAergic interneurons and EC synaptic activity (Cirelli, 2017; Paul *et al.,* 2020; Vyazovskiy *et al.,* 2008).

Recently we reported that constitutive and synaptic GABA_B_/GIRK activity distinguishes mature GCs from adult-born immature GCs (Gonzalez *et al.,* 2018). Here we show a lack of constitutive GABA_B_/GIRK activity in mature GCs during the Dark phase. Most basal activity of GIRK is Gβγ-dependent thus changes in Gβγ availability likely underlie the constitutive GIRK conductance (Rishal et al., 2005). This idea is supported by our observation of a larger constitutive Na^+^ conductance during the Dark phase. This non-selective Na^+^ leak conductance may result from NALCN channels since it is potentiated by low extracellular Ca^2+^ and inhibited by Gβγ (Lu *et al.,* 2010; Philippart and Khaliq, 2018). Thus, the presence of free Gβγ during the Light phase both activates a GIRK conductance while inhibiting NALCN channels, but other conductances could also be affected. Diurnal variation in free Gβγ could result from several mechanisms. Regulators of G-protein signaling (RGS) bind to activated Ga subunits to accelerate GTPase activity thereby reducing free Gβγ (Lin et al., 2014), and both activators of G-protein signaling (AGS) and G-protein receptors kinases (GRK) linked to arrestins can regulate Gβγ availability (Gurevich and Gurevich, 2019). Many components that regulate the GTP signaling system are circadian regulated genes, suggesting that G-protein regulated membrane excitability may also be a conserved mechanism in other populations of neurons across different organisms (Doi *et al.,* 2011; Flourakis *et al.,* 2015; Nakagawa et al., 2020; Noya *et al.,* 2019; Olmedo et al., 2012; Takahashi et al., 2003).

We linked the anti-phase regulation of K^+^ and Na^+^ channels to the circadian molecular clock using conditional deletion of *Bmal1*, a core clock gene. Remarkably, *Bmal1* deletion caused GCs during the Light phase to phenocopy the higher excitability of the Dark phase, with no apparent differences during the Dark phase. In addition to robust but transient expression in developing DG, activation of *Pomc* promoter elements during brain development can generate Cre-mediated recombination in distributed neuronal populations, including the hypothalamic nuclei and SCN (Overstreet et al., 2004). However, our experimental design argues against a major involvement of other cell populations in the effects on GC excitability. First, cKO mice exhibit normal circadian rhythms in both LD and DD conditions. Second, during the Light phase, neighboring *Bmal1* cKO and WT GCs differ in intrinsic properties and afferent recruitment, excluding the possibility that a diffusible substance or other network component mediates the difference in excitability. Importantly, we did not detect differences between tdT^−^ GCs in *Bmal1^lox/lox^* mice and GCs in WT mice, or between tdT^+^ and tdT^−^ GCs in *Pomc-Cre:tdT:Bmal1^wt/wt^*, further ruling out non-specific effects of transient Cre expression. However, potential discordance between recombination of loxp allelles likely underestimates the differences between tdT^+^ and tdT^−^ GCs in *Pomc-Cre:tdT:Bmal1^lox/lox^* mice (Dause and Kirby, 2020).

While the effects of *Bmal1* deletion on GC intrinsic excitability can thus be attributed to cell-autonomous regulation, we cannot rule out a contribution of additional cell populations or mechanisms to the differences we found in DG *c-fos* activation and behavior. While open field activity and the freezing response to strong fear conditioning were unaltered in cKO mice, we found an increase in c-Fos^+^ GCs after exposure to the fear context and improvement in a spatial learning task that was previously shown to be dependent on both the DG excitability and time of day (Pofahl *et al.,* 2021; Snider *et al.,* 2016). These findings agree with other studies showing conditional disruption of the molecular clock in forebrain neurons independently of SCN alter memory consolidation and memory retrieval (Hasegawa et al., 2019; Shimizu et al., 2016; Snider *et al.,* 2016) But in contrast to deletion of *Bmal1* using CamKII-Cre that impaired performance of a similar OLM task during the Dark (Snider *et al.,* 2016), we found improved performance during the Light when mice typically perform poorly. This difference is likely due to selective targeting of *Bmal1* deletion to DG, and is in accordance with the recent finding that optogenetic suppression of GC excitability impairs performance on this type of task (Pofahl *et al.,* 2021).

In conclusion, our results provide new insight into the role of the local molecular clock in the control of neuronal excitability. In addition to the antiphase regulation of passive resting membrane conductances, we also observed changes in the pattern of GC spiking that suggests additional voltage-gated properties are controlled by diurnal regulation of G-protein signaling. Short, high frequency bursts of GCs spikes have been reported in active GCs *in vivo*, and there is increased propensity for this form of bursting in GCs from epileptic tissue (Kelly and Beck, 2017; Pernia-Andrade and Jonas, 2014). Considering the established role of the DG as the hippocampal gate for seizure propagation and recent evidence that disruption of circadian transcriptional loops modify seizure thresholds (Wu *et al.,* 2021; Zhang *et al.,* 2021), our findings present a promising inroad to address a chronotherapeutic approach in the management of TLE (Karoly et al., 2021).

## Acknowledgements

This work was supported by American Epilepsy Society fellowship (JCG); NIH R01NS082413 (KLG), NIH R01NS113948 (JIW) and R01NS064025 & R01NS105438 (LOW). We thank all members of the Wadiche labs for helpful comments throughout this project, and Angela Hill and Mary Seelig for technical assistance.

## Author contribution

Conceptualization and Methodology, J.C.G. and L.O-W.; Formal Analysis and Investigation, J.C.G, H. L., A.M.V. L.K.G, K.L.G and L.O-W.; Resources, G.D.K., K.L.G., J.I.W. and L.O-W.; Writing-Original Draft, J.C.G. and L.O-W.; Writing-Review and Editing, J.C.G., L.K.G., G.D.K., K.L.G., J.I.W and L.O-W.; Supervision and Project Administration, G.D.K., K.L.G. J.I.W. and L.O-W.; Funding Acquisition, J.C.G., K.L.G. J.I.W. and L.O-W.

## STAR METHODS

### RESOURCE AVAILABILITY

#### Lead Contact

Further information and request for resources and reagents should be directed to and will be fulfilled by the lead contact, Linda Overstreet-Wadiche.

#### Materials availability

This study did not generate new unique plasmids, mouse lines or reagents.

#### Data and code availability

- All data reported in this paper will be shared by the lead contact upon request.
- This paper does not report original code.
- Any additional information required to reanalyze the data reported in this paper is available from the lead contact upon request.

### EXPERIMENTAL MODEL AND SUBJECT DETAILS

#### Animals

We used two to five month-old male and female mice from colonies of WT C57BL/6J (Jackson #000664), nNOS-CreERt2 (Jackson #014541); PV-Cre (Jackson #017320); Ai14 (Jackson #007914) and Ai32 (Jackson #024109) mice, all maintained on a C57BL/6J background. Diurnal differences in GC intrinsic properties were detected in all mouse lines (**Figure S1**), so data were pooled. To conditionally delete *Bmal1* from dentate GCs, first we crossed *Pomc-Cre* mice (Jackson # 010714; on a C57Bl/6J background) with *Bmal1^lox^* mice (Jackson # 007668) to generate *Pomc-Cre^+^:Bmal1^lox/lox^* mice. These were subsequently crossed with Ai14 reporter mice (Jackson #007914) to obtain *Pomc-Cre^+^:Bmal1^lox/lox^:tdT*^+^ (*Bmal1* cKO) mice used in experiments (**Figure S6A**). As controls, we also crossed *Pomc-Cre* mice with Ai14 reporter mice (*Pomc-Cre*^+^:*Bmal1*^+/+^:*tdT*^+^). Genotypes were identified by PCR using following primer sets: for Cre: 12048 (CTT GGC TTG GAG GTC TTC TG); 12049 (ACA TGG GTC TCA GAG GCA AT); oIMR 1084 (GCG GTC TGG CAG TAA AAA CTA TC) and oIMR 1085 (GTG AAA CAG CAT TGC TGT CAC TT). For *Bmal1*: oIMR 7525 (ACT GGA AGT AAC TTT ATC AAA CTG) and oIMR 7526 (CTG ACC AAC TTG CTA ACA ATT A). Mice were housed in standard cages with ad libitum access to food and water and maintained in 12:12 light/dark (LD) cycle, except animals used for actograms and recordings in constant darkness (DD). These mice were housed individually in cages equipped with running wheels (Coulbourn Instruments, Whitehall, PA, USA) in LD for at least 15 days before being released into DD. During constant darkness, Circadian time (CT) 12 equals activity onset and the beginning of subjective night. All procedures were approved by the University of Alabama at Birmingham Institutional Animal Care and use Committee (IACUC) in accordance with the US National Institute of Health Guide for the Care and Use of Laboratory Animals.

### METHODS DETAILS

#### Slice preparation

Slices were prepared from mice sacrificed at ZT 5.5 or ZT 11.5 for Light and Dark phase recordings, respectively. Mice were anesthetized with 4% Isoflurane (Fluriso™, USP; Vet One), and 2,2,2-tribromoethanol (Avertin; Sigma-Aldrich) and perfused intracardially with ice-cold modified artificial cerebrospinal fluid (ACSF) cutting solution containing the following (in mM): 110 choline chloride, 7 MgCl_2_, 2.5 KCl, 1.25 Na_2_PO_4_, 0.5 CaCl_2_, 1.3 Na-ascorbate, 3 Na-pyruvate, 25 D-glucose and 25 NaHCO_3_, bubbled with 5% CO_2_ / 95% O_2_. The brain was quickly removed, and 300 μm horizontal slices were prepared using a vibratome (VT1200S, Leica instruments). Slices were incubated at 37°C for ~30 min, then transferred to room temperature in recording solution containing the following (in mM): 125 NaCl, 2.5 KCl, 1.25 Na_2_PO_4_, 1 MgCl_2_, 2 CaCl_2_, 25 D-glucose and 25 NaHCO_3_ bubbled with 5% CO_2_ / 95% O_2_. Slices were also prepared at ZT 0 and ZT 18 for Fig. 2. ZT 18 and DD mice were handled under dim light and perfused with the eyes covered to avoid light interference. Recordings were started two hours after slice preparation during a 3-hour window (ZT 8-11, Light phase or CT 8-11 Subjective Day; ZT 14-17, Dark phase or CT 14-17, Subjective Night), except for Figure 2, where additional cells were recorded before and after the recording window to cover the entire 24 hours cycle.

#### Electrophysiology

Visually identified granule cells in the suprapyramidal and infrapyramidal blade were patched in the whole-cell configuration. GCs expressing tdTomato were visualized using epifluorescence illumination and a Texas Red filter set. GCs in the middle of the granule cell layer were targeted for recording to avoid immature adult-born GCs and semilunar GCs. GCs with intrinsic properties consist with immature or semilunar GCs were discarded. Fire-polished borosilicate glass electrodes (BF150-86-10, Sutter Instrument) with resistance of 2-4 MΩ when filled with intracellular solution were mounted on the headstage (CV-7B) of a Multiclamp 700A amplifier (Molecular Devices). Pipettes for current clamp recordings were filled with the following (in mM): 135 K-gluconate, 3 KCl, 2 MgCl_2_, 10 HEPES, 0.1 EGTA, 10 phosphocreatine, 2 Mg-ATP and 0.5 Na-GTP (excluded in GTP^−^ experiments), pH 7.3 and 310 mOsm. After cancellation of pipette capacitive transients, intrinsic properties were measured in current-clamp mode (no injected current) after using bridge balance to compensate series resistance (< 25mΩ). Voltages were not corrected for junction potentials, calculated to be 13.6 mV. All recordings were made at 30°C and experiments were discarded if substantial changes bridge balance were detected. Currents were sampled at 10kHz and filtered at 2kHz (Digidata 1440A; Molecular Devices) using PClamp 10 software (Molecular Devices). Stimulation of the perforant path (100 ms; 20-100 in increments of 10 mA and 200 mA, 3-5 sweeps per intensity at 10 s intervals) was achieved using a patch pipette (1-2 MΩ) filled with recording solution and controlled by a constant-current isolated stimulator (Digitimer Ltd). To avoid directly stimulating local GABAergic interneurons, electrodes were placed at a distance from the recorded cells, adjacent to the hippocampal fissure between the suprapyramidal blade and apex of the GCL (defined as the midpoint between the suprapyramidal and infrapyramidal blades) to stimulate both lateral and medial perforant path. Sequential “paired” recordings of *Bmal1* cKO and neighboring Bmal1WT GCs were made without changing the stimulation parameters.

Current steps (500 ms) were injected in increments of 10 pA from −10 to 150 pA, and the number of spikes were quantified for each step to generate an input-output curve of neuronal excitability. Rheobase was defined by the minimum current needed to elicit an action potential. Intrinsic passive properties including RMP, IR and TC were calculated using the average of 50 sweeps from a −10 pA current injection.

For voltage clamp experiments, pipettes were filled with (in mM): 122 CsMeSO_3_, 1.8 MgCl_2_, 9 HEPES, 0.45 EGTA, 14 phosphocreatine, 0.09 CaCl_2_, 4 Mg-ATP and 0.5 Na-GTP. Recording solution was composed of (in mM): 125 NaCl, 2.5 KCl, 1MgCl_2_, 1.25 Na_2_PO_4_, 2 CaCl_2_, 25 NaHCO_3_ and 25 D-glucose (bubbled with 95% O_2_ / 5% CO_2_). For Na^+^ substitution, 125 NaCl was replaced with 125 N-Methyl-D-glutamine (NMDG) (Philippart and Khaliq, 2018). GCs were held at −70 mV and recordings were performed in presence of external NBQX (10 μM), R-CPP (50 μM), PTX (50 μM), TTX (1 μM), BaCl_2_ (100 μM), apamin (300 nM) and CsCl_2_ (3 mM).

Data were analyzed with Clampfit (Molecular Devices) and Axograph X (Axograph Scientific). Drugs and chemicals were obtained from Ascent Scientific, Sigma-Aldrich or Tocris Bioscience.

#### Immunohistochemistry

Mice were intracardially perfused with 0.1 M PBS followed by chilled 4% paraformaldehyde (PFA) before brains were removed and post-fixed overnight in 4% PFA. Fifty micrometer free-floating horizontal sections through the entire brain (Vibratome 3000) were stored at −20°C in antifreeze solution (30% ethylene glycol, 20% glycerol in PBS). For BMAL1 immunostaining, slices were washed in PBS and permeabilized and blocked with 0.3% Triton X-100 and 5% BSA for one hour at room temperature. Then sections were incubated overnight at 4°C with anti-BMAL1 antibody (1:2000; NB100-2288, rabbit anti-BMAL1; Novus Biologicals), washed and incubated with goat anti-rabbit Alexa Fluor 488 (1:1000; Invitrogen; 1 hour at room temperature). Sections were mounted using Vectashield without DAPI and imaged using an Olympus FV1200 Confocal Microscope on a 40x water-immersion objective. Images from *Bmal1* cKO and controls were acquired with the same microscope settings and analyzed without adjustment using ImageJ software using a linear region of interest (ROI) drawn across the soma of 3-5 neighboring tdT^+^ and tdT^−^ cells. The Plot Profile option was used to obtain an arbitrary gray value along each point of the ROI in both the red and green channels, and the R value for the correlation at each point was found using Prism (GraphPad Software).

For c-Fos immunostaining slices were washed in TBST (50 mM Tris, 0.9% NaCl and 0.5% Triton X-100) and blocked with TBST + 10% normal goat serum followed by 24-hour incubation at room temperature in guinea pig anti-c-Fos polyclonal antibody (1:1500; Synaptic Systems). This was followed by 2 hours incubation with goat anti-guinea pig Alexa Fluor 488 (1:1000; Invitrogen). Slices were mounted with Prolong Gold mounting medium (Invitrogen). Confocal images were taken on an Olympus Fluoview 1200 with a 40x water-immersion objective. c-Fos+ cells were counted using a BZ-X700 all-in-one Fluorescence Microscope (Keyence, IL, USA).

#### Behavior

Behavior analyses were performed during Light window (ZT 8-11). Researchers conducting and analyzing behavior assays were blind to genotype. Mice were habituated to researchers for 7 consecutive days prior to testing sequentially through open field, object location memory (OLM) and context-dependent fear conditioning tasks (for details see (Laszczyk et al., 2017)). Briefly, for open field assessment of basic movement, mice were placed in the center of an open field apparatus (43 × 43 × 30 cm plexiglass box) with photo beam detectors (ENV-515 software, Med Associates, St. Albans, VT, USA) to quantify activity over 5 minutes. Some cohorts of mice were placed in empty OLM plexiglass testing chamber with activity monitored by Ethovision system XT (Noldus, VA, USA). For OLM testing, mice were habituated to a white plexiglass testing chamber (39 × 19 × 21 cm) with fresh bedding material for 2 consecutive days. On training day, 2 similar plastic bath toy objects were placed on one side of the box and mice were allowed to freely explore for 10 minutes(Haettig et al., 2011). Twenty-four hours later, mice were returned to the box for 5 minutes where 1 object was displaced to the center of the box (see Figure 8E for schematic). Mouse behavior was recorded (TopScan, Clever Sys 2.0, Reston, VA, USA) and videos were manually scored by a different blinded investigator. Interaction was judged to occur if mice were sniffing within 2 cm of a given object. Mice had to explore each object for at least 5 seconds during the training phase and 15 seconds during testing to be included; no mice failed minimum interaction requirements. During training, mice were excluded if they displayed object/side preference greater than 20%; 1 mouse required exclusion. Percent discrimination was calculated as % of time spent with the moved object–time spent with the non-moved object/total time(Wang et al., 2014). For context-dependent fear conditioning, mice were placed in an operant chamber inside an isolation box (Med Associates) for 5 minutes. Mice freely explored for the first 2 minutes and then a series of 3 shocks (1s, 0.5 mA) were delivered (1 x minute). Mice remained in the chamber for 2 minutes after the last shock. 24 hours after training, mice were tested by return to the same chamber for 5 minutes. All training and testing were recorded by automated video tracking system (Med Associates). Percent of time spent freezing was manually scored by measuring freezing behavior in 5 second intervals.

For actograms mice were individually housed with running wheels initially in a 12:12 light-dark cycle (LD) for at least 15 days before being released into constant dark (DD). Wheel-running activity was recorded and analyzed using ClockLab software (Actimetrics, Wilmette, IL). Behavior was analyzed across 10 days of activity. Activity levels were calculated using the batch analysis function in ClockLab software.

### QUANTIFICATION AND STATISTICAL ANALYSIS

Data is presented as median and quartile lines in violin plots and as mean ± s.e.m. in dot plots and histograms. To compare number of spikes elicited by current steps, areas under the curve were measured as suggested by statistic guide of GraphPad followed by a t test or one-way ANOVA. Group comparison used chi-squared (χ^2^), two-sample paired or unpaired t-test or ANOVA with data sets that satisfied normality criteria, while non-normal data sets were analyzed with Mann-Whitney test (Prism; GraphPad Software). For actograms, the free-running period and amplitude were determined by chi-squared (χ^2^) periodogram analysis. Data were analyzed using a one-way ANOVA with LSD post-hoc tests. Rhythmic comparison of intrinsic properties was calculated using Cosinor analysis using SPSS (version 25) and the following equation:

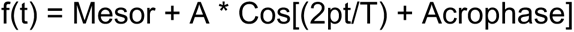

where

Mesor (acronym for middle estimating statistic of rhythm) = the mean of the oscillation
A = the amplitude (peak-to-trough difference)
T = the period (24 h)
Acrophase = the timing of the cosine maximum
t = a timepoint

R^2^ is the resulting statistic that measures the percent variance accounted for by the 24 h approximating waveform.

