## Supplemental material for "Circadian regulation of dentate gyrus excitability mediated by G-protein signaling"

### SUPLEMENTARY FIGURE 1

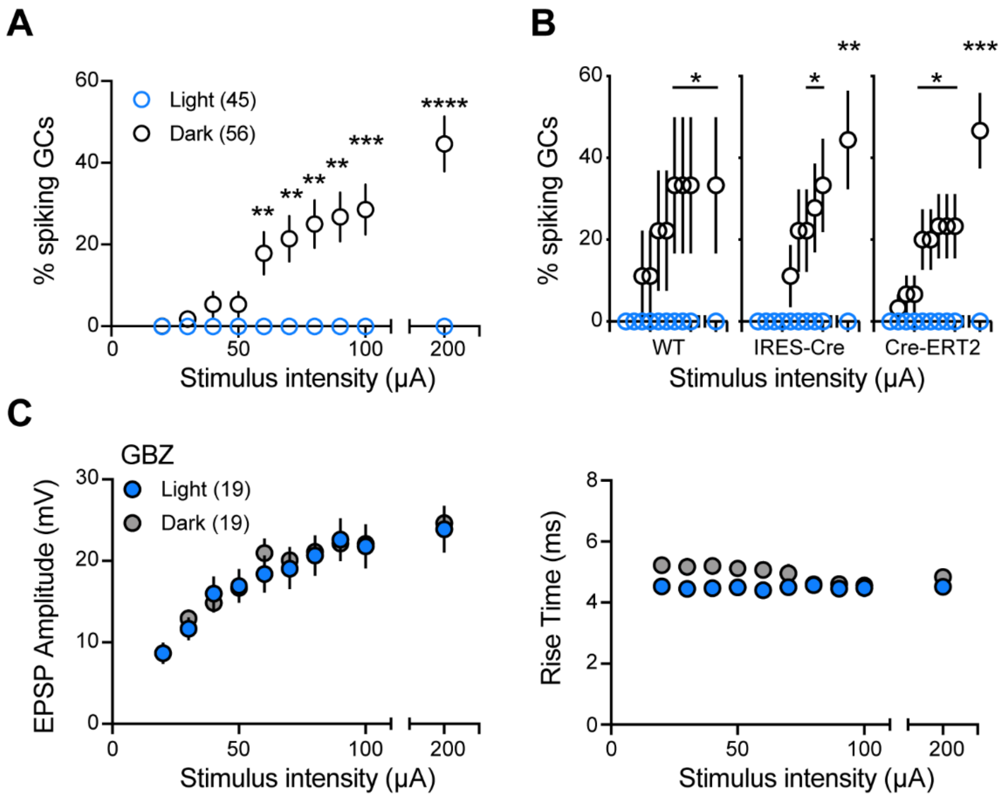


**Figure S1, related to Figure 1. Diurnal differences in recruitment of GC spiking is robust across mouse lines and additional analysis of subthreshold EPSPs.**

(A) Percentage of GCs spiking in response to perforant path stimulation at increasing stimulus intensities, showing results pooled across multiple mouse lines (shown in B) and including the experiments shown in Fig 1b. n = 45, 56 for Light and Dark.

(B) Results separated by transgenic mouse lines. Sample sizes: wild type (WT C57Bl/6) n = 10, 9; constitutive Cre, PV-Cre crossed with Ai32 (IRES-Cre) n = 14, 18 and inducible Cre, nNOS-CreERt2 crossed with Ai32 (Cre-ERT2) n = 21, 29 for Light (blue) and Dark (black) respectively.

(A-B) $\chi^2$ tests, *p<0.05; **p<0.01; ***p<0.001; ****p<0.0001.

(C) Additional analysis of subthreshold EPSPs from Figure 1d showing that neither the amplitude of EPSPs measured from a normalized RMP (right) nor the 20-80% rise time of EPSPs (right) differed between the Light and Dark phase. Experiments in GBZ. Multiple comparison test p>0.05, n = 19,19.

#### SUPLEMENTARY FIGURE 2

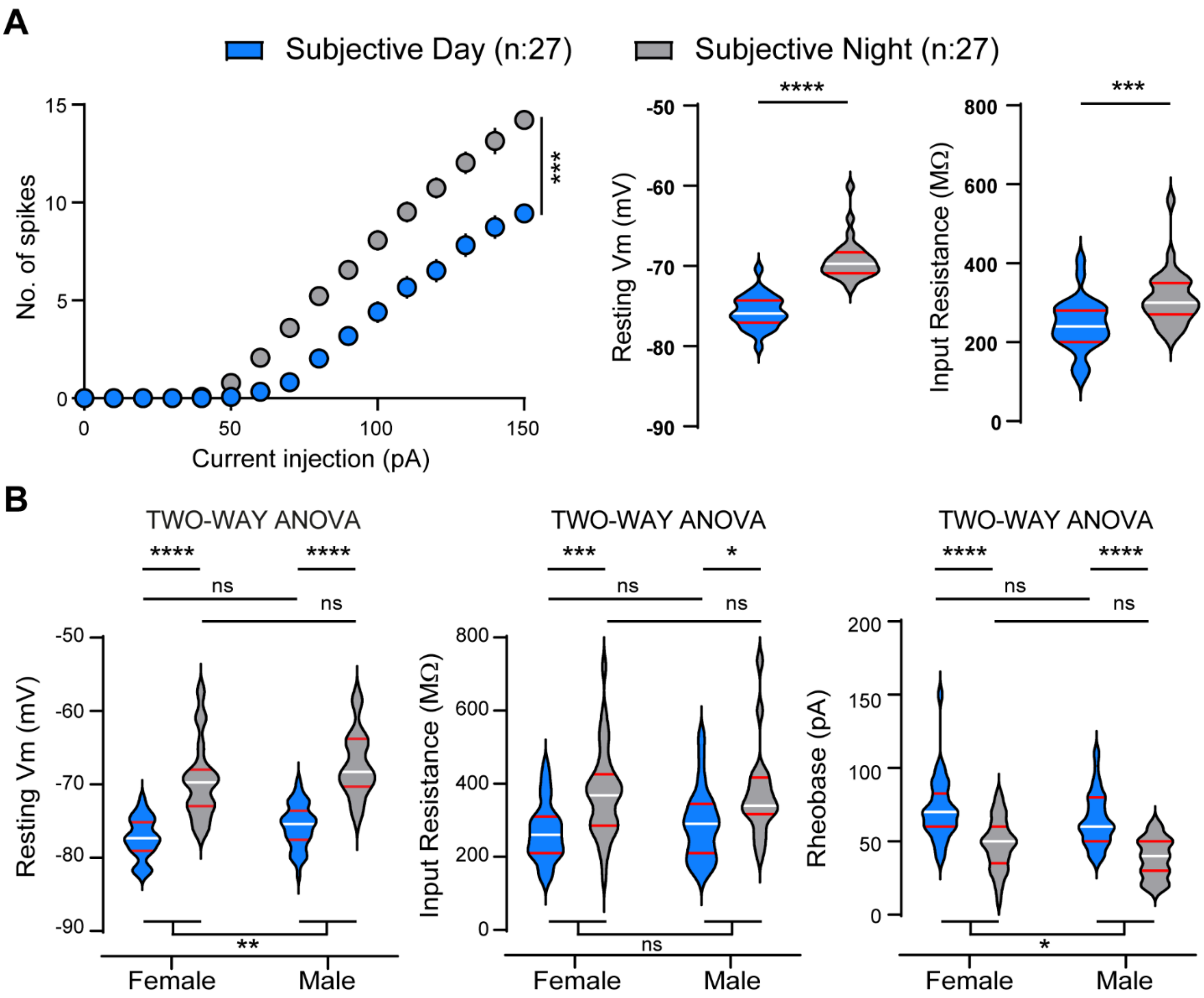


**Figure S2, related to Figure 2. Differences in GC intrinsic excitability persist in constant darkness.**

(A) Number of spikes elicited by increasing current steps, RMP and IR from GCs in mice housed in constant darkness. Recordings were performed in male mice individually housed in activity cages, with Subjective Day (circadian time; CT 8-11) and Subjective Night (CT 14-17) phases identified by initiation of locomotor activity at CT12. Unpaired t-test to compare area under the curve (AUC) t = 4 ***$p<0.001$. Unpaired t-test, t = 10.3 and t = 3.6 for RMP and IR respectively, *** $p < 0.001$; **** $p < 0.0001$.

**(B)** Violin plots showing intrinsic properties of GCs from male and female mice recorded during Light and Dark phase (as in Figure 1). Two-way ANOVA showed no interaction between sex and recording time (see Table S2 for details).

#### SUPLEMENTARY FIGURE 3

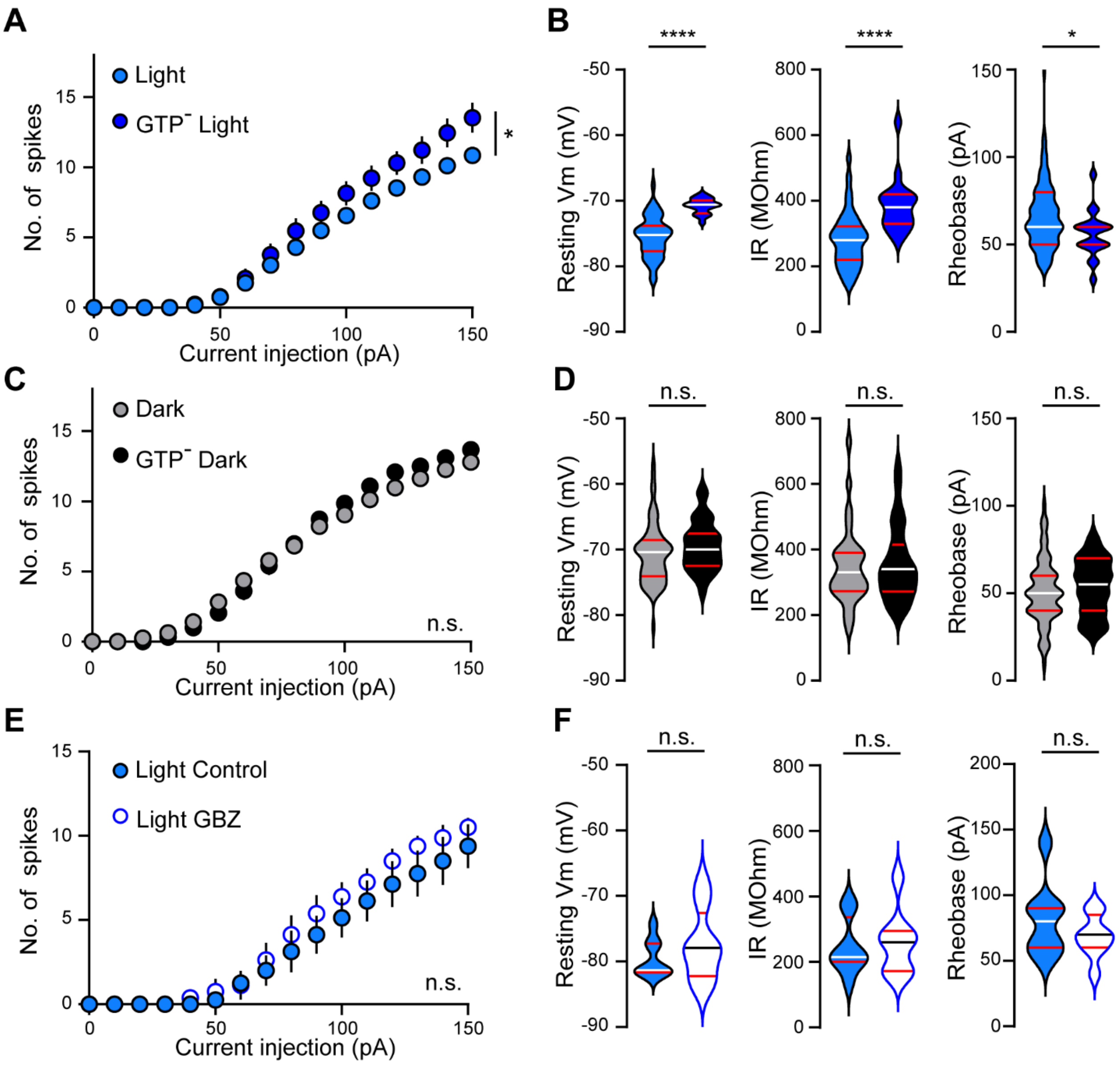


**Figure S3, related to Figure 3. Intracellular GTP affects intrinsic properties only during the Light phase.**

(A, C, E) Number of spikes elicited by increasing current steps and (B, D, E) violin plots showing intrinsic properties in GCs recorded with or without Na-GTP (GTP⁻) in the intracellular solution (A-D) or in presence of GBZ 10 μM (E-F).

(A-B) In GCs during the Light phase, GTP⁻ increased intrinsic excitability. Welch's ANOVA to compare area under the curve (AUC) F = 2.6, Mann-Whitney U test, U = 124.5; unpaired t-test, t = 4.9 and t = 2.2 for RMP, IR and rheobase respectively. * p < 0.05; **** p < 0.0001.

(C-D) During the Dark phase, GTP did not affect intrinsic excitability. Welch's ANOVA to compare area under the curve (AUC) $F = 0.4$, Mann-Whitney U test, $U = 1.3$; unpaired t-test, $t = 0.4$ and $t = 0.7$ for RMP, IR and rheobase respectively. $p>0.05$.

(E-F) GCs excitability is not altered in presence of GBZ. Welch's ANOVA to compare area under the curve (AUC) $F = 1.4$, paired t-test $t = 1$ $t = 0.4$ and $t = 1.2$ for RMP, IR and rheobase respectively. $p>0.05$.

#### SUPLEMENTARY FIGURE 4

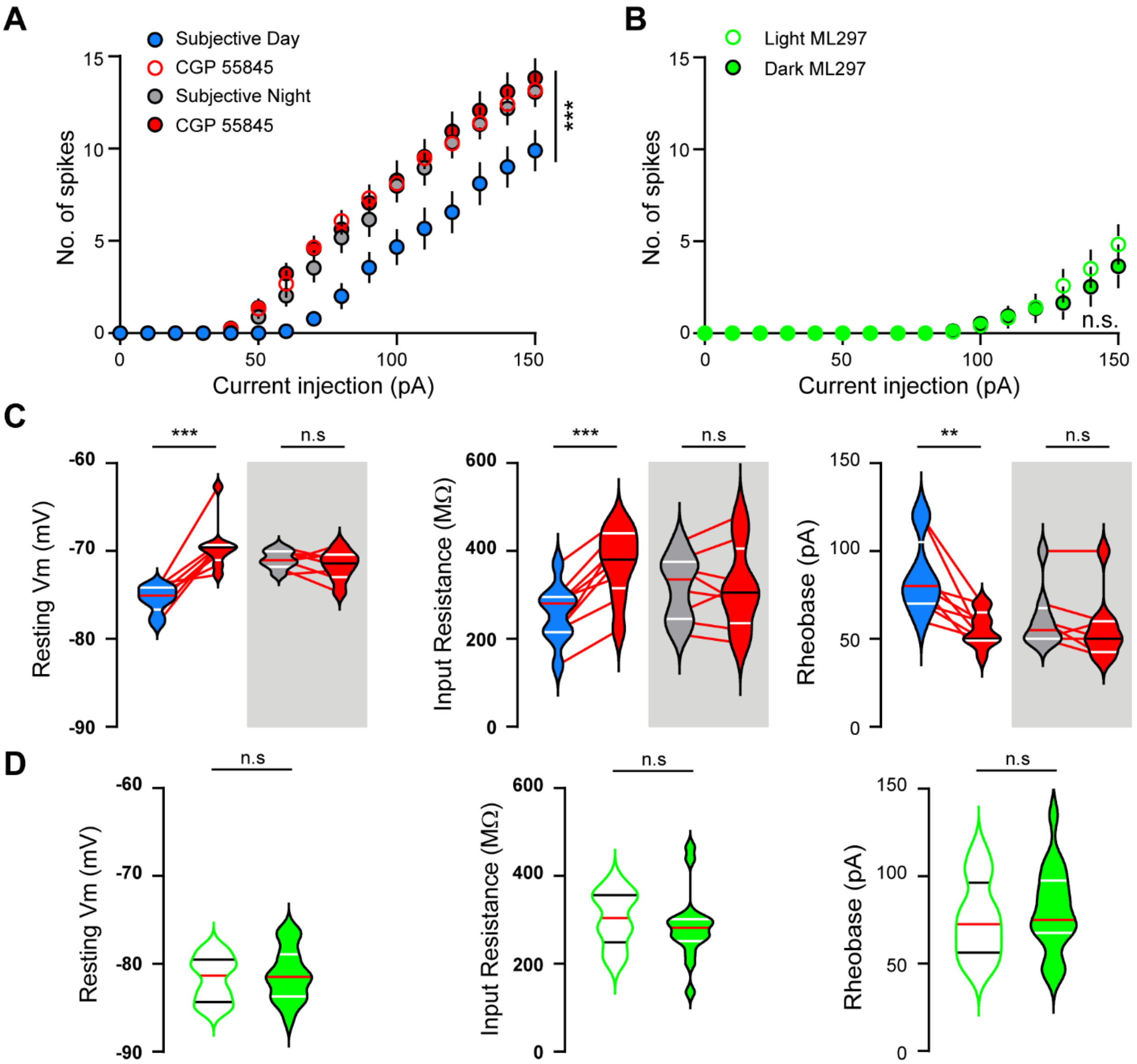


**Figure S4, related to Figure 4. Differential effects of CGP persist in constant darkness.**

(A, C) Number of spikes elicited by current steps and violin plots showing intrinsic properties recorded from male mice housed in DD. CGP enhances intrinsic excitability only during Subjective Light but not Subjective Dark phases. One-way ANOVA to compare area under the curve (AUC) $F_{(3,30)} = 9.5$ followed by Fisher LSD comparisons test. ***p<0.001. Paired t-test t = 5.8; t = 6.6 and t = 4.2 for RMP, IR and rheobase respectively. ** p < 0.01; *** p < 0.001.

(B, D) Additional analysis of data in Figure 4b showing that the GIRK activator ML297 normalizes GCs properties across Light and Dark phases. Unpaired t-test to compare area under the curve (AUC) t = 0.2. Unpaired t-test t = 0.4, t = 0.6 and t = 0.5 for RMP, IR and rheobase respectively. p>0.05.

#### SUPLEMENTARY FIGURE 5

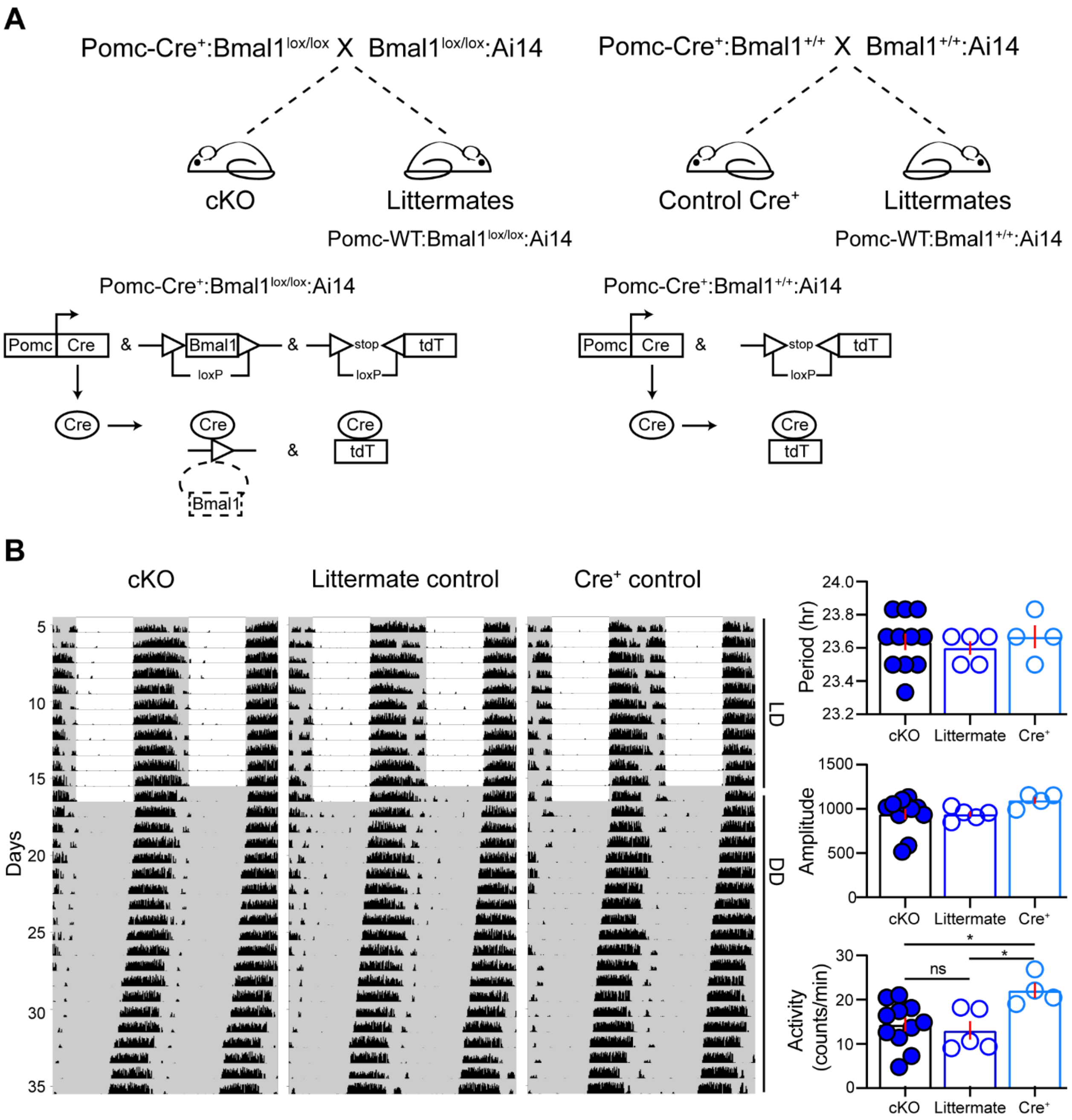


**Figure S5, related to Figure 6. Breeding strategy for *Bmal1* cKO mice that exhibit normal circadian behavior.**

(A) Schematic illustrating conditional excision of the *lox-p* flanked *Bmal1* locus using proopiomelanocortin (*Pomc*) promoter. Cre activity eliminates the *Bmal1* locus and expresses tdTomato (tdT). Cre negative littermates and Cre+ (*Pomc-Cre*: Ai14) were used as controls for actograms. Cre+ (*Pomc-Cre*: Ai14) mice were used as controls for recordings and behavior. No differences in litter size, gender or Cre ratio were observed when comparing cKO with *Pomc-Cre*:tdT controls.

(B) Representative double-plotted actograms of running wheel activity (left) and quantification of behavioral rhythms and activity levels in DD (right). Cre+ mice had slight increase in wheel activity counts, but no difference in open field behavior. See Table 3 and 4 for additional details.

**SUPLEMENTARY FIGURE 6**

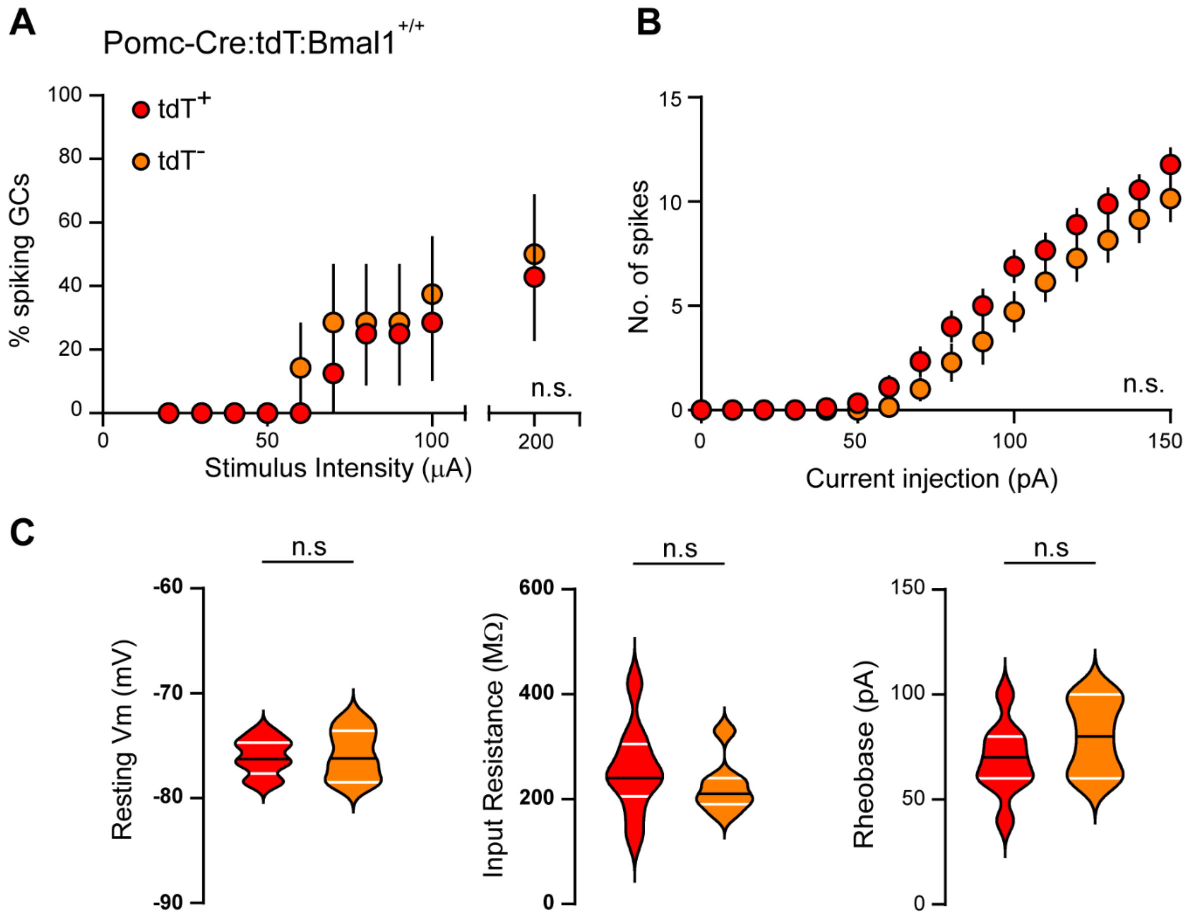


**Figure S6, related to Figure 7. No differences in synaptic recruitment or intrinsic excitability of tdT+ and tdT- GCs in control mice.**

(A) Percentage of GCs spiking in response to perforant path stimulation during the Light phase, comparing $tdT^+$ and tdT- GCs in *Pomc-Cre:tdT* control mice as shown in Figs 6 & 7. $\chi^2$ tests, p>0.05.

(B) Number of spikes elicited by current injection and

(C) intrinsic properties from GCs $tdT^+$ and $tdT^-$. Unpaired t-test to compare area under the curve (AUC) t = 2. Unpaired t-test t = 0.2, t = 0.9 and t = 1.2 for RMP, IR and rheobase respectively n = 9, 7 for $tdT^+$ and $tdT^-$.

**Supplemental Table S1, related to Figure 2.**

Cosinor analysis of intrinsic properties of GCs (estimate ± SE)

| | Rhythmicity | | Cosinor Output Parameters | | |
|---|---|---|---|---|---|
| Intrinsic properties | $R^2$ | P Value | Mesor | Amplitude | Acrophase |
| RMP | 0.375 | P<0.0001 | -71.8 ± 0.2 | 3.77 ± 0.27 | 21.2 ± 0.3 |
| Input Resistance | 0.168 | P<0.0001 | 287.2 ± 5.1 | 60.2 ± 7.6 | 15.9 ± 0.4 |
| Time Constant | 0.280 | P<0.0001 | 26.9 ± 0.4 | 7.5 ± 0.7 | 15.0 ± 0.3 |
| Rheobase | 0.167 | P<0.0001 | 63.8 ± 1.0 | 11.9 ± 1.5 | 4.3 ± 0.5 |

**Supplemental Table S2, related to Figure S2.**

Two-factor ANOVA of Intrinsic properties of GCs

| | Sex | | Circadian | | Interaction | |
|---|---|---|---|---|---|---|
| Intrinsic properties | F | P Value | F | P Value | F | P Value |
| RMP | $F_{(1,127)}$=7.1 | P<0.01 | $F_{(1,127)}$=131.1 | P<0.0001 | $F_{(1,127)}$=0.08 | 0.76 |
| Input Resistance | $F_{(1,127)}$=0.26 | 0.60 | $F_{(1,127)}$=24.6 | P<0.0001 | $F_{(1,127)}$=0.34 | 0.55 |
| Time Constant | $F_{(1,127)}$=0.03 | 0.86 | $F_{(1,127)}$=24.8 | P<0.0001 | $F_{(1,127)}$=0.73 | 0.39 |
| Rheobase | $F_{(1,127)}$=6.57 | P<0.05 | $F_{(1,127)}$=55.7 | P<0.0001 | $F_{(1,127)}$=0.06 | 0.79 |

**Supplemental Table S3, related to Figure S5.**

Summary (mean ± SEM) of circadian behavioral parameters and ANOVA results in LD.

| | **cKO**<br>*n=11* | **littermate**<br>*n=5* | **Cre+ controls**<br>*n=4* | **ANOVA**<br>*F (P)* |
|---|---|---|---|---|
| Activity (revolutions/min) | 13.7 ± 1.6 | 13.14 ±1.57 | 20.18 ± 0.96 | 3.379 (0.058) |
| Light Activity (revolutions/day) | 607.2 ± 351.8 | 1060.24 ± 590.9 | 940.5 ± 467.1 | 0.301 (0.744) |
| Dark Activity (revolutions/day) | 19202.6 ± 2435.6 | 17862.2 ± 1942.6 | 28119.2 ± 1429.2 | 3.220 (0.065) |
| Total Activity (revolutions/day) | 19809.9 ± 2356.9 | 18922.4 ± 2271.1 | 29059.7 ± 1393.7 | 3.379 (0.058) |
| Percent Light Activity (%) | 3.73 ± 2.46 | 4.97 ± 2.28 | 3.22 ± 1.62 | 0.083 (0.921) |

**Supplemental Table S4, related to Figure S5.**

Summary (mean ± SEM) of circadian behavioral parameters and ANOVA results in DD.

| | **cKO** *n=11* | **littermate** *n=5* | **Cre+ controls** *n=4* | **ANOVA** *F (P)* |
|---|---|---|---|---|
| Period (hour) | 23.6 ± 0.04 | 23.6 ± 0.04 | 23.66 ± 0.06 | 0.240 (0.789) |
| Period Amplitude | 940.2 ± 61.1 | 940.5 ± 30.4 | 1098.4 ± 38.8 | 1.517 (0.248) |
| Activity (revolutions/min) | 14.37 ± 1.54 | 13.03 ± 2.07 | 22.1 ± 1.7 | 4.837 (0.022) |
